# Genetic diversity unveiled: cost-effective methods for grassland species

**DOI:** 10.1101/2025.07.23.666374

**Authors:** Damian Käch, Miguel Loera-Sánchez, Bruno Studer, Roland Kölliker

## Abstract

Permanent grasslands are predominantly composed of allogamous plant species that exhibit high levels of plant genetic diversity (PGD) within their populations. Grasslands with high PGD are more resilient to environmental stress and constitute valuable reservoirs of genetic resources for plant breeding. Therefore, monitoring PGD is the basis for detecting changes in PGD and correspondingly intervening. However, PGD monitoring is often neglected in biodiversity reports due to difficulties in taking representative samples and in using standardised and affordable indicators of PGD. Here we successfully applied two common approaches, multispecies amplicon sequencing (MSAS) and genotyping-by-sequencing (GBS), to assess PGD of agronomically relevant grassland species. Using MSAS, we were able to taxonomically distinguish five species (*Dactylis glomerata* L., *Festuca pratensis* Huds., *Lolium perenne* L., *Trifolium pratense* L., *Trifolium repens* L.) from multispecies samples and differentiate accessions within species, with fixation index (*F* _ST_) values ranging from 0.014 for *T. repens* to 0.089 for *L. perenne*. Based on an extended *L. perenne* sample set containing mixtures of two cultivars at different ratios, mixtures containing both cultivars at 50% separated from the corresponding cultivars according to this ratio using MSAS and GBS. Furthermore, GBS enabled separation of samples containing two cultivars at a 75:25-ratio from the corresponding cultivars and the 50:50-ratio samples. These results indicate complementing applications of the two approaches in PGD monitoring. While we anticipate that MSAS with its cost-effectiveness could be applied to large-scale PGD monitoring, GBS with its lower detection limit could be applied to studies where cultivar composition shifts are of interest.

## 1 Introduction

Permanent grasslands are fundamental to sustainable ruminant livestock production and provide multiple ecosystem services (Schils et al., 2022). They cover 42.8% of the agricultural area of Western Europe and, in some countries, this proportion reaches over 60% (e.g., Ireland, the United Kingdom or Switzerland (Boch et al., 2020; Lüscher et al., 2019)). These grasslands are composed of economically important plant species, most of which are obligate outcrossers from the Poaceae (grass) and Fabaceae (legume) families, resulting in populations with high genetic diversity (i.e., intraspecific diversity) (Last et al., 2013). This plant genetic diversity (PGD) is essential for the functioning and resilience of grassland ecosystems. It plays a crucial role in providing ecosystem services, such as reducing invertebrate herbivory (Wan et al., 2022) and stabilising biomass productivity (Meilhac et al., 2019; Prieto et al., 2015). Additionally, grasslands of high PGD constitute valuable reservoirs of genetic resources for forage plant breeding. The prospect of leveraging these benefits has led to initiatives aimed at fostering the *in situ* protection and management of PGD in grasslands.

Monitoring PGD in permanent grasslands is key to detect undesired changes, intervene accordingly, and thus protect these valuable ecosystems. However, these shifts cannot be detected visually and require molecular genetic approaches. Advances in DNA sequencing have enabled the study of PGD in many non-model species with complex and large genomes, including grassland plant species, at continuously decreasing costs (Loera-Sánchez et al., 2019). Sequencing-based genetic diversity assessment is usually conducted based on single-nucleotide polymorphism (SNP) data, which is used to produce the SNP allele frequencies needed to estimate common genetic diversity and differentiation metrics like nucleotide diversity, heterozygosity and fixation indices (*F* _ST_). A diverse array of genetic approaches is available that generate SNP data covering several targeted regions to whole genomes. These approaches enable the study of tens to hundreds of populations or accessions, as has been shown for some of the major plant species of permanent grasslands (Nay et al., 2023; Faville et al., 2020; Tamura et al., 2023).

Amplicon sequencing and genotyping-by-sequencing (GBS) are two common approaches to assess PGD in grassland species. For amplicon sequencing, a method was recently proposed as a low-resolution (10^2^ to 10^3^ SNPs), cost-effective approach for PGD assessment (Loera-Sánchez et al., 2022). This multispecies amplicon sequencing (MSAS) method is based on the targeted sequencing of single-copy, nuclear orthologous genes and ultra-conserved-like elements (ULEs) present in multiple forage grass and legume species, using MSAS primers that are transferable across multiple species within each plant family. GBS, on the other hand, uses restriction enzymes for genome complexity reduction and generates SNP data at higher resolution (10^4^ to 10^5^ SNPs) covering loci across the whole genome (Elshire et al., 2011). Therefore, MSAS and GBS differ mainly in the amount and distribution of loci that their SNP data cover, suggesting that each method could be suitable for different purposes in PGD monitoring.

Despite the availability of various molecular-genetic methods and the decreasing cost of DNA sequencing, genetic diversity monitoring is often neglected in national biodiversity conservation reports worldwide (Hoban et al., 2023). This has been attributed to a lack of standardized and affordable indicators of genetic diversity in natural populations (Hoban et al., 2021b). In the context of PGD monitoring in grassland, this could be in part due to logistical challenges to produce representative samples from large land extensions, which can harbor a great diversity of plant communities due to variations in microtopography. Grasslands, which commonly consist of multiple species, each represented by various accessions, pose additional questions regarding the representation of each species and accession. These complexities extend to the subsequent steps of the workflow, where appropriate genetic and bioinformatics methods have to be selected depending on the aim of PGD monitoring, whether it is detecting shifts in species or even accession composition.

In this study, evaluate the suitability of MSAS for efficient PGD monitoring in species from permanent grasslands. Our specific objectives are to: (i) assess the genetic differentiation in pairs of accessions from five major species (*Dactylis glomerata* L., *Festuca pratensis* Huds., *Lolium perenne* L., *Trifolium pratense* L., and *Trifolium repens* L.) using MSAS; (ii) test the suitability of MSAS to distinguish cultivars and detect cultivar shifts; and (iii) compare these results to that of GBS using an extended sample set of *L. perenne*. Based on these analysis, we discuss the prospects for long-term, multispecies PGD monitoring in grasslands.

## 2 Materials and Methods

We used MSAS to measure genetic differentiation in pairs of accessions of five major species from permanent grasslands (*D. glomerata*, *F. pratensis*, *L. perenne*, *T. pratense*, and *T. repens*). We further focused on six *L. perenne* cultivars to assess the detection limits and variability of genetic differentiation (i.e., fixation index, *F* _ST_) and structure metrics based on MSAS by comparing them to those produced with a genome-wide approach (GBS).

### 2.1 Plant material and DNA extraction

#### 2.1.1 Seedling pools

For the single-accession seedling samples (SA samples), seeds of *D. glomerata*, *F. pratensis*, *L. perenne*, *T. pratense*, and *T. repens* were germinated on filter paper in Petri dishes. Two accessions per species were used. We will use the label ”A” for the first accession of each species, and ”B” for the second (Table S1). For each accession, DNA extraction was performed on three biological replicates (pool of 20 seedlings per replicate). The grass seedlings were sampled at approximately 2 cm from the tip, discarding the radicule. The legume seedlings were sampled when the cotyledons emerged, discarding the radicule. The plant material was stored for a maximum of two weeks at -70°C in 1.5 ml tubes until processed. They were subsequently ground in 1.5 ml tubes using plastic pestles while keeping the tubes half-submerged in liquid nitrogen. DNA was extracted from the ground material using the NucleoSpin Plant II kit (Macherey-Nagel, Duren, Germany) resulting in 30 SA samples (five species, two accessions per species in three biological replicates).

Additionally, mixed-species samples were prepared by pooling DNA from the SA samples. The mixed-species seedling samples (MS samples) were prepared in triplicate for three different compositions: 1) MS-A100, for equal DNA amounts of accessions A, 2) MS-B100, for equal amounts accessions B, and 3) MS-AB50, for equal amounts of MS-A100 and MS-B100 (Table S2). Biological replicates of SA samples were kept separate for the preparation of MS samples resulting in six MS samples.

#### 2.1.2 Extended *L. perenne* sampling

We chose perennial ryegrass to conduct a wider sampling design, considering single plants and swards. For establishing the single plants, seeds of six perennial ryegrass cultivars (‘Arara‘, ‘Araias‘, ‘Repentinia‘, ‘Artonis‘, ‘Arcturus‘, ‘Algira‘) were germinated on filter paper in Petri dishes. All cultivars are early heading and were bred and multiplied by Agroscope (Zurich, Switzerland) and Delley seeds and plants Ltd (Dellay, Switzerland; Suter et al. 2025, Suter et al. 2017). While ‘Arara‘, ‘Araias‘, and ‘Repentinia‘ are diploid (2n = 2x = 14), ‘Artonis‘, ‘Arcturus‘, and ‘Algira‘ are tetraploid (2n = 4x = 28) cultivars. After germination, the seedlings were transferred into pot trays (88 wells, 37×27 cm, compost as substrate). Pooled-plant samples were prepared and consisted of one or two cultivars (Tab. 1). Single-cultivar samples contained 30 plants represented by one leaf fragment of 5 cm per plant. For each of these cultivar compositions, three biological replicates were prepared (i.e., three replicates with 30 different plants per replicate). In addition to these pure samples, mixtures of two cultivars were prepared. These mixed samples contained a total of 60 plants, i.e., 30 plants of ‘Araias‘ and 30 plants of another cultivar, at two mixing ratios. Samples with a 50:50-ratio contained leaf fragments of 5 cm with one leaf fragment per plant. Samples with a 75:25-ratio contained leaf fragments of 7.5 cm and 2.5 cm of ‘Araias‘ and another cultivar, respectively, with one leaf fragment per plant.

**Table 1:**
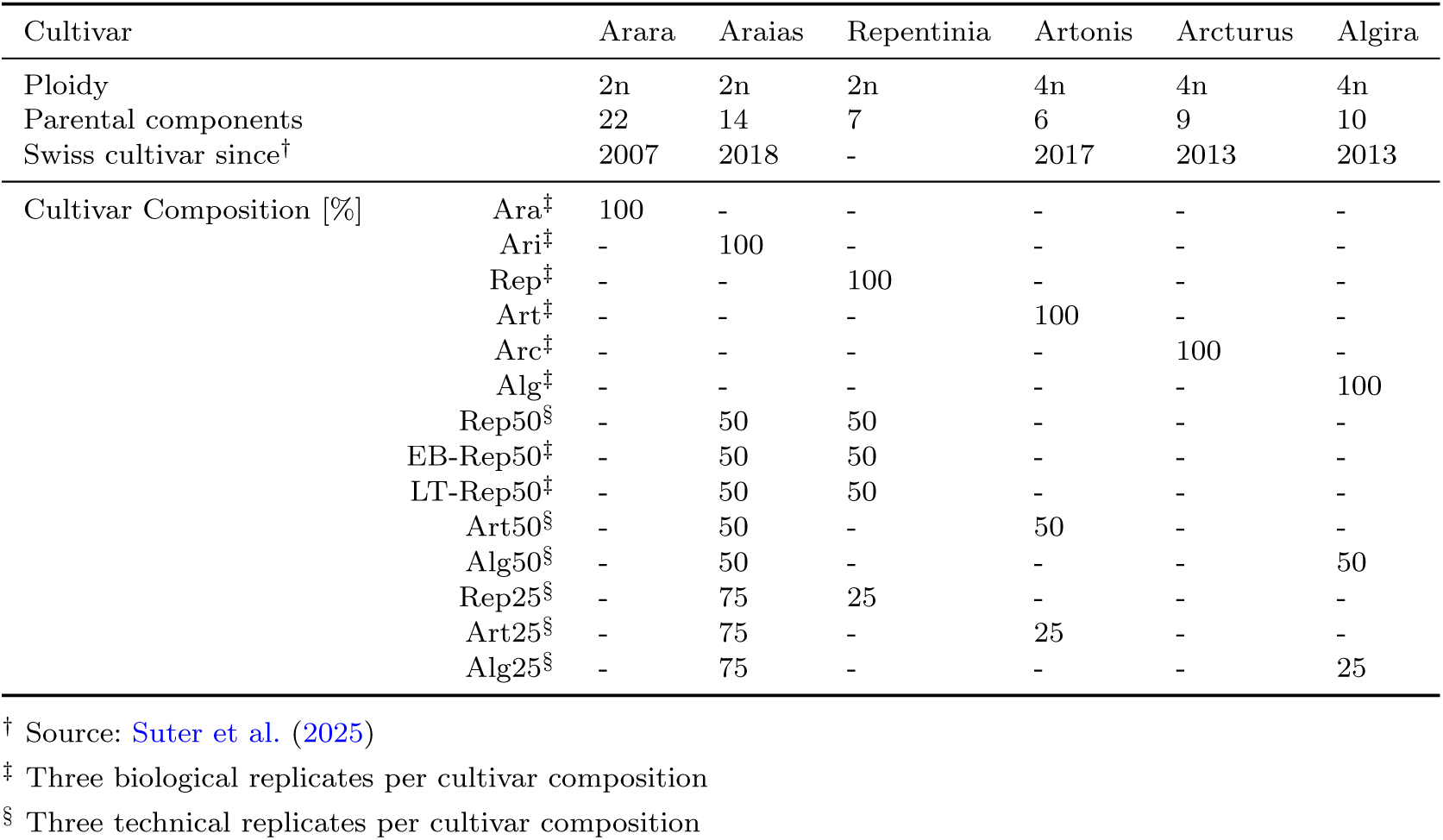
Cultivar compositions based on six early heading *Lolium perenne* L. cultivars bred and multiplied by Agroscope (Zurich, Switzerland) and Delley seeds and plants Ltd (Delley, Switzerland), respectively, and grown in greenhouse and field experiments at Eschenbach (EB) and Langenthal (LT).

In addition to the single plants in the greenhouse experiment, seeds of ‘Araias‘ and ‘Repentinia‘ were mixed at a 50:50-ratio and sown in field plots of 1.5 by 6 m at a sowing density of 500 seeds/m^2^. These cultivar compositions were grown in three replicates at two locations in Switzerland (Langenthal BE [47*^◦^*13*^′^*3.40*^′′^*N, 7*^◦^*48*^′^*15.24*^′′^*E] (LT), Eschenbach LU [47*^◦^*8*^′^*16.54*^′′^*N, 8*^◦^*18*^′^*16.89*^′′^*E] (EB)). After sward establishment, the plants were sampled along six parallel transects per plot. Evenly distributed along each transect, 30 plants were sampled by taking four leaf fragments of ∼5 cm length per plant. For each plot, two of the six transects were randomly selected and the leaf fragments were cut to a length of 1 cm resulting in 60 plants-pools per plot. The samples from both the greenhouse and the field experiment were stored at -80°C, freeze-dried for 72 hours, and disrupted in a Qiagen TissueLyser II (Qiagen, Hilden, Germany).

DNA was extracted using the NucleoSpin II kit (Macherey-Nagel, Düren, Germany) and its purity and concentration was assessed using a NanoDrop spectrophotometer (Thermo Fisher Scientific, Waltham, MA, USA). Based on these measurements, DNA was normalised to a concentration of 30 ng/µl. Technical replicates of the mixed samples from the greenhouse experiment were generated by dividing the normalised DNA into three samples resulting in a total of 42 samples.

### 2.2 MSAS

Nine previously reported (Loera-Sánchez et al., 2022) and 22 newly designed multispecies primers for grass and legume species were used for MSAS (Table 2). Primers were designed based on data from Loera-Sánchez et al. (2022) and they target portions of single-copy orthologous genes and ultra-conserved like elements that display within-species variability in forage grass or legume species.

**Table 2:**
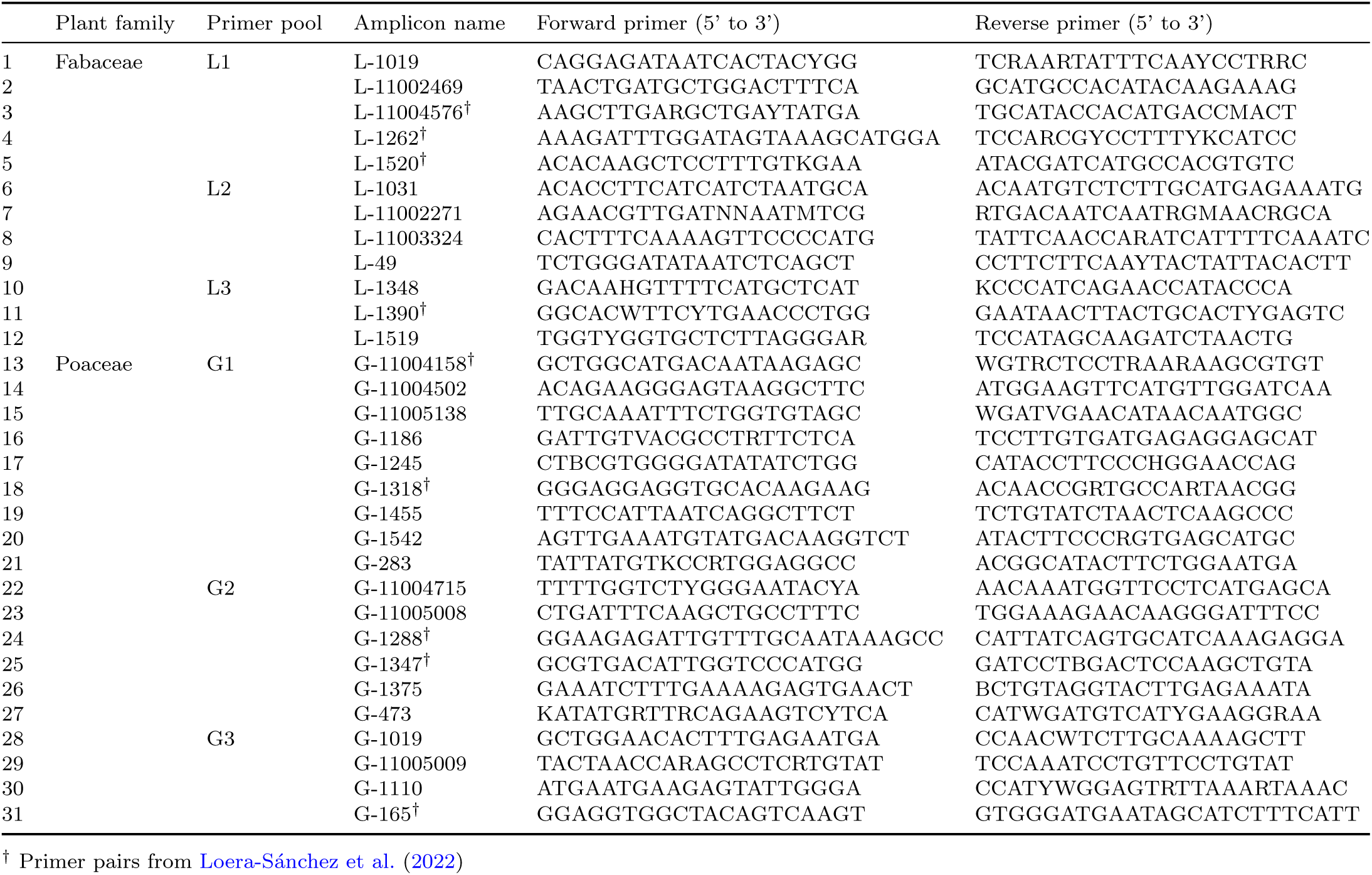
Primer pairs used for multispecies amplicon sequencing.

Multiplex PCR was performed with the seedling samples and with the extended *L. perenne* samples. We used the Qiagen Multiplex PCR kit (Qiagen, Hilden, Germany) with 5 µl Qiagen multiplex PCR buffer, 1 µl Q-solution, 1 µl primer mix (10 µM; three different mixes per sample; Table 2), and 15 ng of DNA in a total volume of 10 µl. In total, we used three primer mixes. The amplification conditions were: 15 minutes at 95°C, 30 seconds at 94°C, then 45 cycles of 2 minutes at 50°C and 1 minute at 72°C. A final extension step ran for 10 min at 72°C, then reactions were held at 10°C. The resulting amplicons were pooled per sample and cleaned using AMPure beads (Beckman Coulter Life Sciences, Brea, CA, USA) using a bead-to-sample ratio 0.9×.

Illumina library preparation was performed separately for the seedling samples and for the *L. perenne* extended sample set using the NEBNext Ultra II kit (New England Biolabs, Ipswich, MA, USA). Dual indexing ran for five cycles using the NEBNext Dual Index Set 1 (New England Biolabs). Samples were then cleaned with AMPure beads (0.9× proportion) and pooled equimolarly. A four-cycle reconditioning PCR was performed for each library pool. Such a reaction consisted of 10 µl water, 12.5 µl KAPA HiFi mix (Roche, Basel, Switzerland), 1 µl i5 primer (10 µM), 1 µl i7 primer (10 µM), and 2.5 µl of the dual-indexed library pool. The thermocycler program was: three minutes at 95°C followed by four cycles of 20 seconds at 98°C, 15 seconds at 62°C, and 30 seconds at 72°C, followed by a final extension step of one minute at 72°C. Library pools were spiked with 10% of phiX and sequenced on an Illumina MiSeq platform (MiSeq reagent kit v3, 2×300bp; Illumina, San Diego, CA, USA) at the Genetic Diversity Center (ETH Zurich, Zurich, Switzerland).

### 2.3 GBS

GBS was conducted by LGC Genomics (Berlin, Germany) using double-enzyme digestion (PstI-MspI). The samples were sequenced on an Illumina NextSeq platform (2 flow cells, 1×75bp; Illumina, San Diego, CA, USA). Only the extended *L. perenne* sample set was processed for GBS.

### 2.4 Data processing and analysis

#### 2.4.1 MSAS variant calling and taxonomical assignment

Amplicon sequencing reads were quality-controlled with fastQC v.0.11.9 (Andrews, 2010) and multiQC v1.9 (Ewels et al., 2016). Quality filtering (Q*>*20), primer and adapter trimming was done using cutadapt v4.3 (Martin, 2011) using the forward and reverse complement of each primer sequence. Reads from the SA samples were mapped to species-specific reference FASTA files with the sequences of the target amplicons using minimap2 v2.17 (Li, 2018). For MS samples, a FASTA file containing the sequences of the target amplicons for 16 grassland plant species was used as a reference (Loera-Sánchez et al., 2022). Reads from the extended *L. perenne* samples were mapped to the *L. perenne* Kyuss reference genome (Chen et al., 2024) using minimap2.

For the SA and the extended *L. perenne* samples, reads that mapped in proper pairs with MAPQ*>*20 were retained using SAMtools v1.15.1 (Danecek et al., 2021) (samtools view -Sb -f 2 -q 20). For MS samples, an additional filtering that retained only primary alignments was performed (samtools view -b -F 256 -q 20). For all sample types, PCR duplicates were marked using Picard v3.0.0 (Broad Institute, 2019) (picard MarkDuplicates) and variant calling was performed using BCFtools v1.15.1 (Danecek et al., 2021) with a maximum number of reads of 2000. Raw variant calls were filtered using BCFtools to retain bi-allelic SNPs with a MAF=1% and QUAL*>*20.

Taxonomic assignment of MS samples was performed at the read mapping stage. This was facilitated by the sequence identifiers in the FASTA reference file, which contained the abbreviation for each species and the amplicon name. Thus, VCF files were split by assigned species.

To estimate taxonomic assignment rates, raw reads from SA samples (i.e., samples for which the species is known) were concatenated by species and then mapped to the 16 species reference FASTA. The number of reads assigned to each reference species was obtained with SAMtools (samtools idxstats). The taxonomic assignment rate was the count of reads correctly assigned to a species divided by the total read count for that species.

#### 2.4.2 GBS variant calling

GBS reads were demultiplexed and adapter / restriction enzyme sequences were removed by the service provider. The pre-processed reads were then quality-controlled and mapped as described above for amplicon sequencing. SAMtools was used to retain only primary alignments and mapped reads (samtools view -Sb -F 260) and to compute coverage (samtools coverage). Variant calling and filtering was conducted with the same parameters like amplicon sequencing. We used GNU parallel v20220522 for read processing and variant calling (Tange, 2011).

#### 2.4.3 Genetic differentiation analysis

Allele frequencies and *F* _ST_ statistics were produced based on filtered VCF files using the R package ’poolfstat’ v2.1.1 (Gautier et al., 2022; Hivert et al., 2018). *F* _ST_ and pairwise *F* _ST_ were calculated for single-accession data (SA, taxonomically-assigned MS except for MS-AB50, and pure extended *L. perenne*). Additionally, for each species, variants of MS samples were merged with those of SA samples, retaining only variants present in both. The resulting dataset (SAMS variants) were also used for *F* _ST_ and pairwise *F* _ST_ calculations.

For all single-accession variant datasets, permutational multivariate analysis of variance (PERMANOVA) ran using the function adonis2 from the R package ’vegan’ v.2.6-2 (Oksanen et al., 2022) on euclidean distance matrices based on SNP allele frequencies, cultivar names as fixed effects, and 10,000 permutations. Discriminant analysis of principal components (DAPC) for SAMS variants was performed based on their allele frequency matrices using the R package ’adegenet’ v2.1.10 (Jombart and Ahmed, 2011; Jombart, 2008), retaining 4 PCA axes (i.e., considering *k* −1 for k=5 *a priori* groups) and 5 axes for the discriminant analysis. For the pure extended *L. perenne* samples, DAPC was performed on allele frequencies. For this, the optimal numbers of PCA axes to be retained were determined by conducting cross-validation using unscaled data and default parameters (i.e., *xvalDapc(…, scale = FALSE)*). The mixed samples were predicted based on that optimised DAPC using default parameters (i.e., *predict.dapc(…)*).

#### 2.4.4 Population structure analysis on the extended *L. perenne* samples

Population structure analysis was performed on the extended *L. perenne* samples by, first, separately calling variants in four groups of samples: 1) the Ari, Rep, Rep50, EB-Rep50, LT-Rep50, and Rep25 samples; 2) the Ari, Art, Art50, and Art25 samples; 3) the Ari, Alg, Alg50, and Alg25 samples; and 4) the pure samples (i.e., Ara, Ari, Rep, Art, Arc, Alg). The cultivar compositions of such samples are shown in Table 1. Each of these four groups of samples was separately analysed using ’STRUCTURE’ v2.3 (Pritchard et al., 2000) and ’ADMIXTURE’ v1.3 (Alexander et al., 2009). For the analysis using ’STRUCTURE’, VCF files were converted to ’STRUCTURE’ format using ’PGDSpider’ v2.1.1.5. (Lischer and Excoffier, 2012) and computed using the parameter settings described in Supporting Information A. For the analysis using ’ADMIXTURE’, the same VCF files were used as for ’STRUCTURE’, were converted to BED format using ’PLINK’ v1.9 (Purcell et al., 2007) and computed using the parameter settings described in Supporting Information A.

Data processing and visualization was conducted using the R packages ’dplyr’ v1.0.10 (Wickham et al., 2022), ’tidyr’ v1.2.1 (Wickham and Girlich, 2022), ’stringr’ v1.5.0 (Wickham, 2022), ’purrr’ v1.0.1 (Wickham and Henry, 2023), ’ggplot2’ v3.4.0 (Wickham, 2009), ’ggrepel’ v0.9.2 (Slowikowski, 2022), and ’ggpubr’ v0.5.0 (Kassambara, 2022). Figures and calculations were performed using R v4.2.3 (R Core Team, 2023) and Rstudio v2023.03.0 Build 386 ”Cherry Blossom” Release (RStudio Team, 2020).

## 3 Results

### 3.1 Sequencing output

The raw MSAS output for the seedling samples was 21.18 million read pairs. After read mapping and filtering, 11.93 million read pairs remained, of which 4.42 million read pairs were from SA samples and 7.52 million read pairs from MS samples.

The raw MSAS output for the extended *L. perenne* sampling was 19.03 million pair-ended reads, which resulted in 12.45 million quality-controlled read pairs after discarding phiX reads (3.39 million read pairs) and other undetermined reads (1.99 million read pairs), removing primer sequences and controlling for read quality and length. Samples Art25-1 and Rep-2 did not produce any sequencing reads and were not considered further. All amplicons were found in the Kyuss reference genome (Tab. 3). 12.37 million read pairs mapped to the reference genome in proper pairs. Across all samples, average coverage per amplicon ranged from 287±415 (ortho-1375) to 133,489±50,571 (ortho-1019).

**Table 3:** Mapping coordinates of the amplicons in the *Lolium perenne* L. Kyuss genome sequence.

| Locus | Chromosome<br>(chr) | Start | End |
| --- | --- | --- | --- |
| G-283 | chr1 | 26882183 | 26882675 |
| G-1542 | chr1 | 142148139 | 142148658 |
| G-1455 | chr2 | 49666450 | 49666893 |
| G-473 | chr3 | 177676768 | 177677250 |
| G-11005138 | chr4 | 271713826 | 271714378 |
| G-1288 | chr4 | 271713865 | 271714449 |
| G-11005009 | chr4 | 354783605 | 354784087 |
| G-11005008 | chr4 | 354784729 | 354785194 |
| G-1375 | chr5 | 28083648 | 28084120 |
| G-165 | chr5 | 207437103 | 207437696 |
| G-11004158 | chr6 | 25898407 | 25898991 |
| G-1186 | chr6 | 108414665 | 108415028 |
| G-11004715 | chr6 | 239751220 | 239751612 |
| G-1347 | chr6 | 242837713 | 242838233 |
| G-11004502 | chr6 | 264461326 | 264461599 |
| G-1019 | chr7 | 53843084 | 53843493 |
| G-1245 | chr7 | 203254074 | 203254613 |
| G-1110 | chr7 | 224883315 | 224883719 |
| G-1318 | chr7 | 310725655 | 310726210 |

The raw GBS output for the extended *L. perenne* sampling was 983.1 million single-end reads. We received 897.6 million quality-controlled reads with removed adapter sequences and restriction enzyme sites. Of these, 607.8 million reads mapped to and, on average, covered 1.7% of the Kyuss reference genome.

### 3.2 Genetic differentiation between accession pairs in five major grassland plant species

For SA samples, the number of variants called ranged from 86 SNPs for *T. pratense*to 802 SNPs for *D. glomerata*. Mean *F* _ST_ values ranged from 0 for *T. repens* to 0.61 for *D. glomerata* (Tab. 4). Between-accessions pairwise *F* _ST_ ranged from 0.006±0.005 for *T. repens* to 0.723±0.006 for *D. glomerata*, while within-accession pairwise *F* _ST_ ranged from -0.001±0.009 for *T. pratense* to 0.009±0.017 for *F. pratensis*. Between- and within-accession pairwise *F* _ST_ values were significantly different for all species except *T. repens* (Wilcoxon rank sum test, *P* ≤ 0.05. Fig. 1(a)). No significant differences in allele frequencies were detected by PERMANOVA (SA in Tab. S3).

**Fig. 1:**
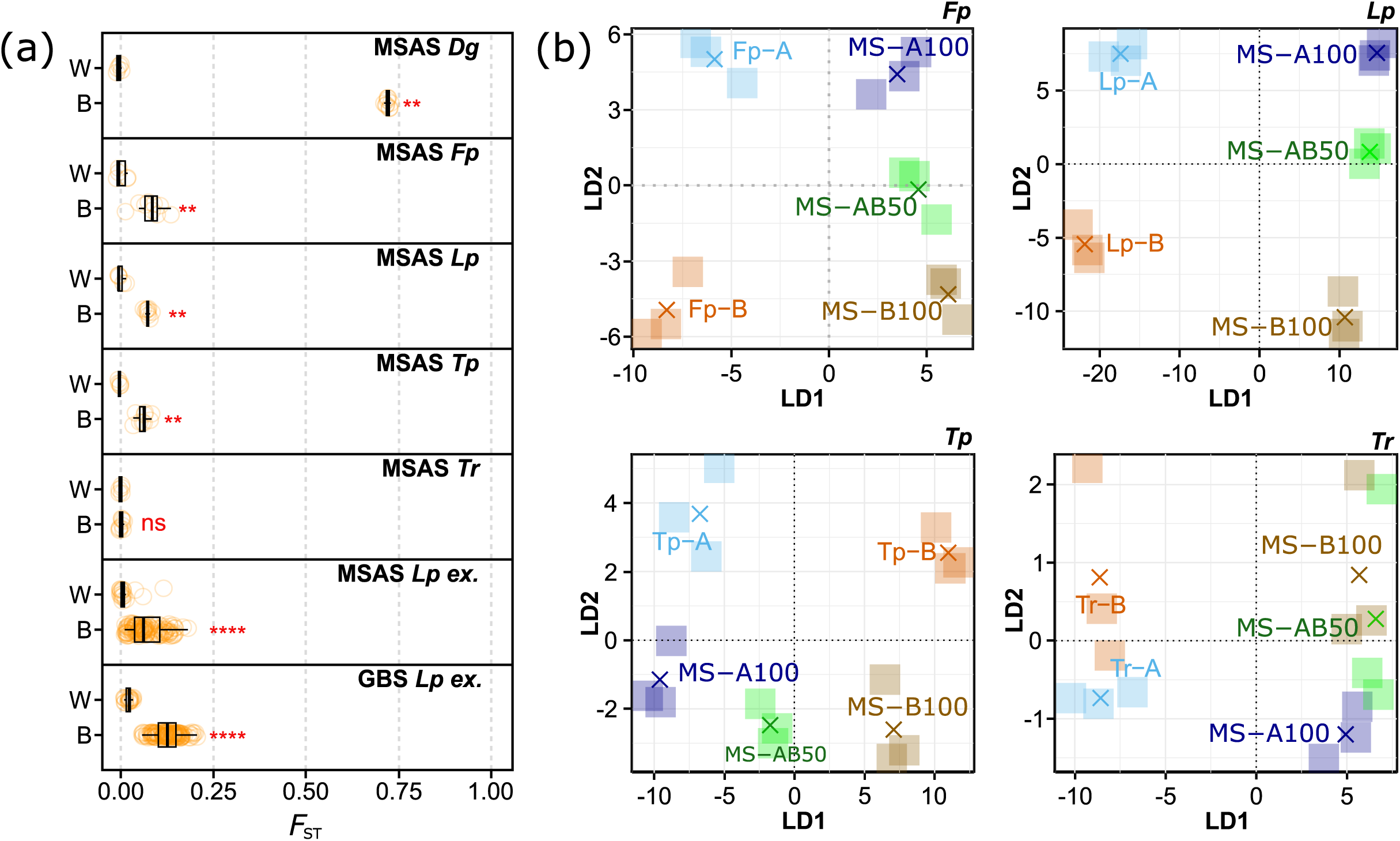
Pairwise within-(W) and between-accession (B) fixation index (*F* _ST_) values based on single-accession seedling samples of five grassland species (*Dactylis glomerata* L. (*Dg*), *Festuca pratensis* Huds. (*Fp*), *Lolium perenne* L. (*Lp*), *Trifolium pratense* L. (*Tp*), *Trifolium repens* L. (*Tr*)) and single-cultivar samples of an extended *L. perenne* sample set (*Lp* ex.; (a)). Datasets were generated using multispecies amplicon sequencing (MSAS) or genotyping-by-sequencing (GBS), indicated in the panel titles. The Wilcoxon rank-sum test significance levels are ns: *P >* 0.05, *: *P* ≤ 0.05, **: *P* ≤ 0.01, ***: *P* ≤ 0.001, ****: *P* ≤ 0.0001. Discriminant analysis of principal components separately for each species based on single-accession seedling samples (two accessions per sample, indicated by A and B), mixtures containing accession A of all species (MS-A100; each accession represented by 20%), mixtures containing accession B of all species (MS-B100; each accession represented by 20%), and 50:50-ratio mixture of MS-A100 and MS-B100 (MS-AB50; (b)). Replicate means are represented by crosses and the dataset was generated using MSAS.

**Table 4:**
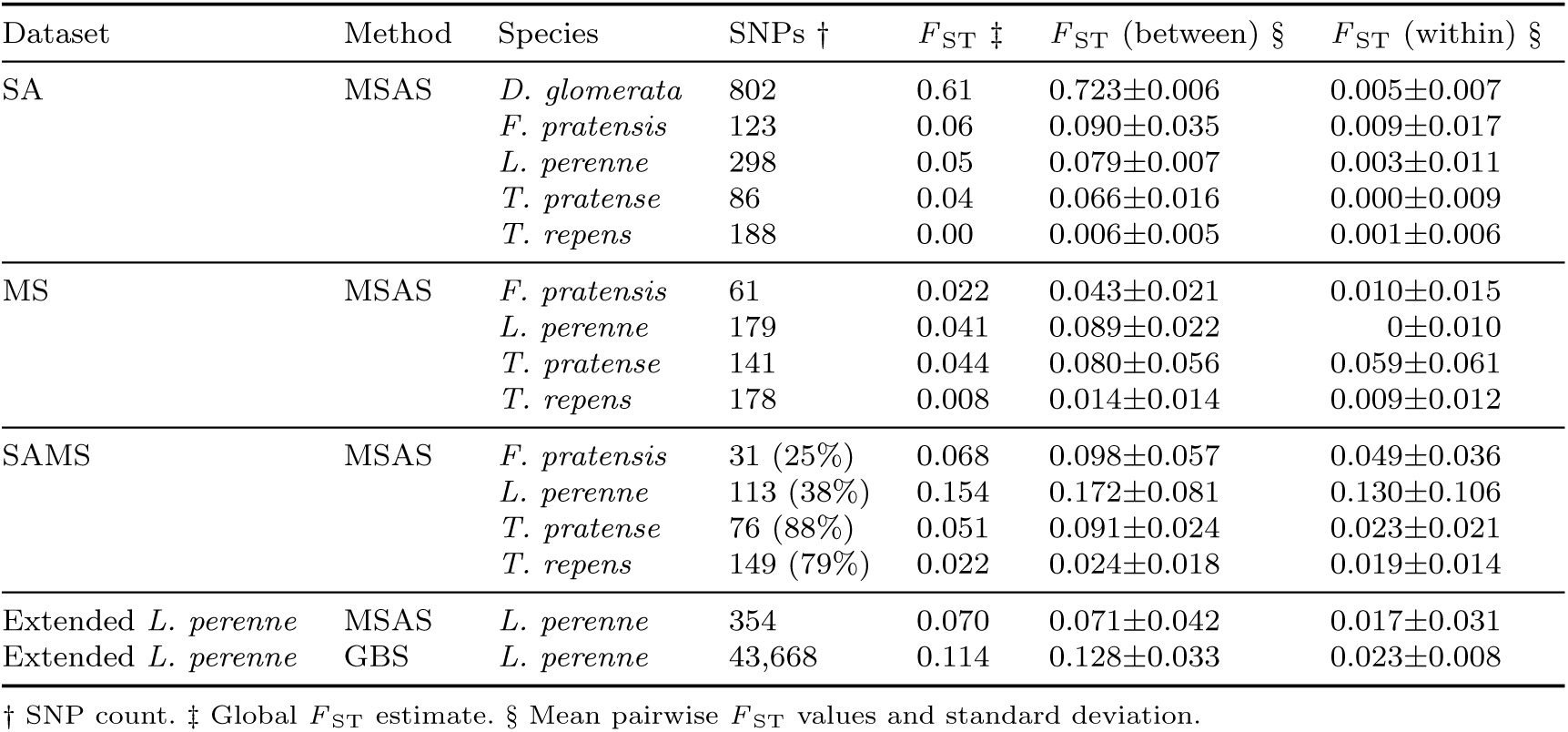
Single-nucleotide polymorphism (SNP) counts, global fixation index (*F* _ST_) estimates, pairwise between- and within-accession *F* _ST_ values for datasets based on single-accession seedling (SA), taxonomically assigned mixed-species seedling (MS), variants present in both SA and MS (SAMS; with percentage of SNPs present in SA), and single-cultivar samples of an extended set of *Lolium perenne* L.. Datasets were generated using multispecies amplicon sequencing (MSAS) or genotyping-by-sequencing (GBS).

### 3.3 Taxonomic assignment and genetic differentiation analysis in mixed-species samples

Correct taxonomic assignment rates were 21.2%, 54.2%, 68.9%, 99.9%, and 73.3% of reads for *D. glomerata*, *F. pratensis*, *L. perenne*, *T. pratense*, and *T. repens*, respectively. Most frequent species incorrectly assigned were *Poa pratensis* L. (38.3% of reads) for *D. glomerata*; *L. perenne* (40.6%) for *F. pratensis*; *Lolium multiflorum* Lam. (13.9%) for *L. perenne*; and *T. pratense* (26.7%) for *T. repens*. Variant calling and filtering of the taxonomically assigned reads resulted in 61, 179, 141, and 178 SNPs in MS samples of *F. pratensis*, *L. perenne*, *T. pratense*, and *T. repens*, respectively (MS in Tab. 4). No SNPs were obtained for taxonomically-assigned, MS *D. glomerata* samples. Overall *F* _ST_ ranged from 0.008 (*T. repens*) to 0.044 (*T. pratense*). Between-accessions pairwise *F* _ST_ ranged from 0.014±0.014 (*T. repens*) to 0.089±0.022 (glsfepra). Within-accessions pairwise *F* _ST_ ranged from 0±0.010 (*F. pratensis*) to 0.059±0.061 (*T. pratense*).

The differentiation pattern observed between accessions from SAMS samples is also in agreement to the pattern observed in SA samples. Variant calling and filtering resulted in 31, 113, 76, and 148 SNPs in SAMS samples *F. pratensis*, *L. perenne*, *T. pratense*, and *T. repens*, respectively (SAMS in Tab. 4). Taxonomic assignment resulted in higher *F* _ST_ estimates for SAMS samples compared to SA samples. Overall *F* _ST_ ranged from 0.022 (*T. repens*) to 0.154 (*L. perenne*). Between-accessions pairwise *F* _ST_ ranged from 0.0.024±0.018 (*T. repens*) to 0.172±0.081 (glsloper). Within-accessions pairwise *F* _ST_ ranged from 0.019±0.014 (*T. repens*) to 0.130±0.106 (*T. pratense*). Significant allele frequency differences were detected for *F. pratensis* and *T. pratense* by PERMANOVA, with 35.20% and 52.90%, respectively, of the total variance attributed to between-accessions variation (SAMS in Tab. S3). The two accessions in each species (except for *T. repens*) and the three types of MS samples were distinguished (Fig. 1(b)). Furthermore, accessions were close to the corresponding MS samples and MS-AB50 samples were located between the MS-A100 and the MS-B100 samples.

### 3.4 Genetic differentiation in an extended sample set of *L. perenne*

For MSAS and GBS, variant calling resulted in 354 and 43,668 SNPs, respectively. Based on pure samples, mean within-cultivar pairwise *F* _ST_ (0.017±0.031 for MSAS and 0.023±0.008 for GBS) was significantly different to between-cultivar pairwise *F* _ST_ (0.071±0.042 for MSAS and 0.128±0.033 for GBS) for both sequencing approaches (Fig. 1(a)).

The cultivars could be differentiated with both GBS and MSAS with mean *F* _ST_ values of 0.114 and 0.070, respectively (Fig. 2, Tab. 4). Between-cultivar *F* _ST_ values of GBS were higher than those of MSAS across all samples. However, the *F* _ST_ values of GBS significantly correlated with those of MSAS (Fig. 2(c)). Between-cultivar variation in GBS allele frequencies amounted to 75.34% of the total variance (PERMANOVA, *P* =0.0001; Table S3). For MSAS allele frequencies, between-cultivar variation was 66.86% of the total variance (PERMANOVA, *P* =0.0001; Table S3).

**Fig. 2:**
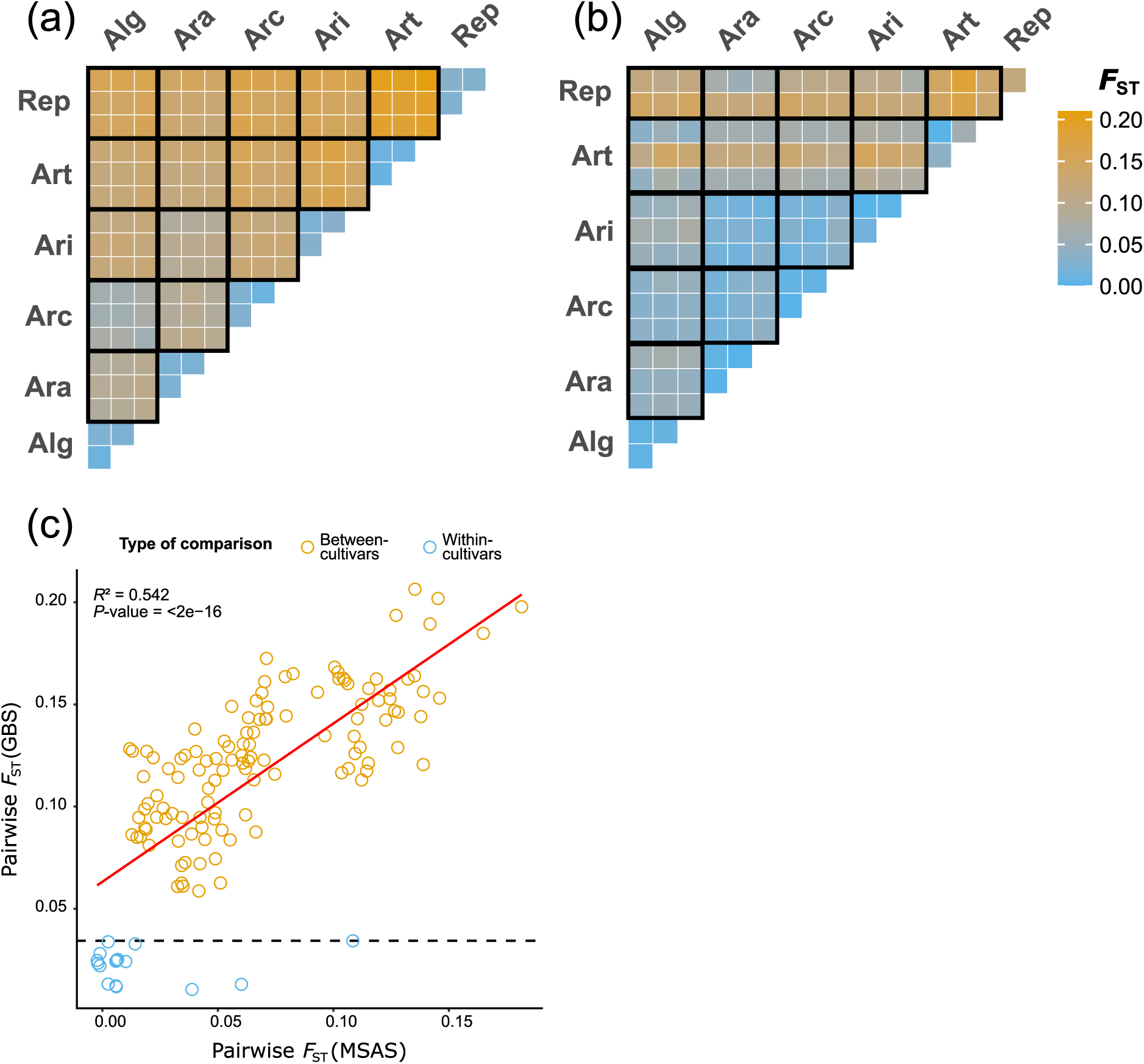
Pairwise fixation index (*F* _ST_) values between the six cultivars ‘Arara‘ (Ara), ‘Araias‘ (Ari), ‘Repentinia‘ (Rep), ‘Artonis‘ (Art), ‘Arcturus‘ (Arc), and ‘Algira‘ (Alg) of an extended *Lolium perenne* L. sample set analysed using genotyping-by-sequencing (GBS;(a)) and multispecies amplicon sequencing (MSAS;(b)). Comparison of pairwise *F* _ST_ values between GBS and MSAS (c), where beige and blue circles refer to between- and within-cultivar comparisons, respectively.

For GBS, *F* _ST_ values between samples of ‘Algira‘ and ‘Arcturus‘ were lower compared to the other cultivars. For MSAS, the samples of ‘Artonis‘ and ‘Repentinia‘ had higher *F* _ST_ compared to the other cultivars. Replicates had the lowest *F* _ST_ values in both sequencing approaches. This high similarity of replicates was also reflected in high correlations of allele frequencies. Correlation coefficients between replicates were larger than 0.90 for both GBS and MSAS (Fig. S1). Allele frequencies different to 0 or 1 were, however, more frequent for GBS than MSAS. Additionally, the similarity of replicates and separation of cultivars could be shown by cluster analysis (Fig. S3, K=6), where replicates were assigned to the same and cultivars to separate clusters. While these differentiation patterns were nearly perfect for GBS (i.e., replicates were fully assigned to the same cluster), some samples of MSAS were partially assigned to multiple clusters. Contrary to these results derived by the ’ADMIXTURE’ analysis, cultivars could not be separated using ’STRUCTURE’: The six cultivars were assigned to four and to one cluster for GBS and MSAS, respectively (Fig. S7, K=6). Nevertheless, the similarity of replicates could be shown by similar cluster assignments using ’STRUCTURE’.

The cultivars could be clearly separated using a DAPC on the GBS data (Fig. 3(a)). The DAPC was based on three PCA axes as this number resulted in the lowest mean square error during the cross-validation procedure. Furthermore, the DAPC separated the cultivar mixtures and the corresponding cultivars according to the mixing ratios and could. The 50:50-ratio samples were located in the middle of the corresponding cultivars. Additionally, the 75:25-ratio samples, which consist 75% of ‘Araias‘ and 25% of another cultivar, were located between the corresponding 50:50-ratio samples and ‘Araias‘. Therefore, the differentiation patterns represent the genetic structure within the samples. These differentiation patterns could also be observed when separating the linear discriminants (LD). However, the differentiation pattern of LD1 is closer to the expected pattern than LD2.

**Fig. 3:**
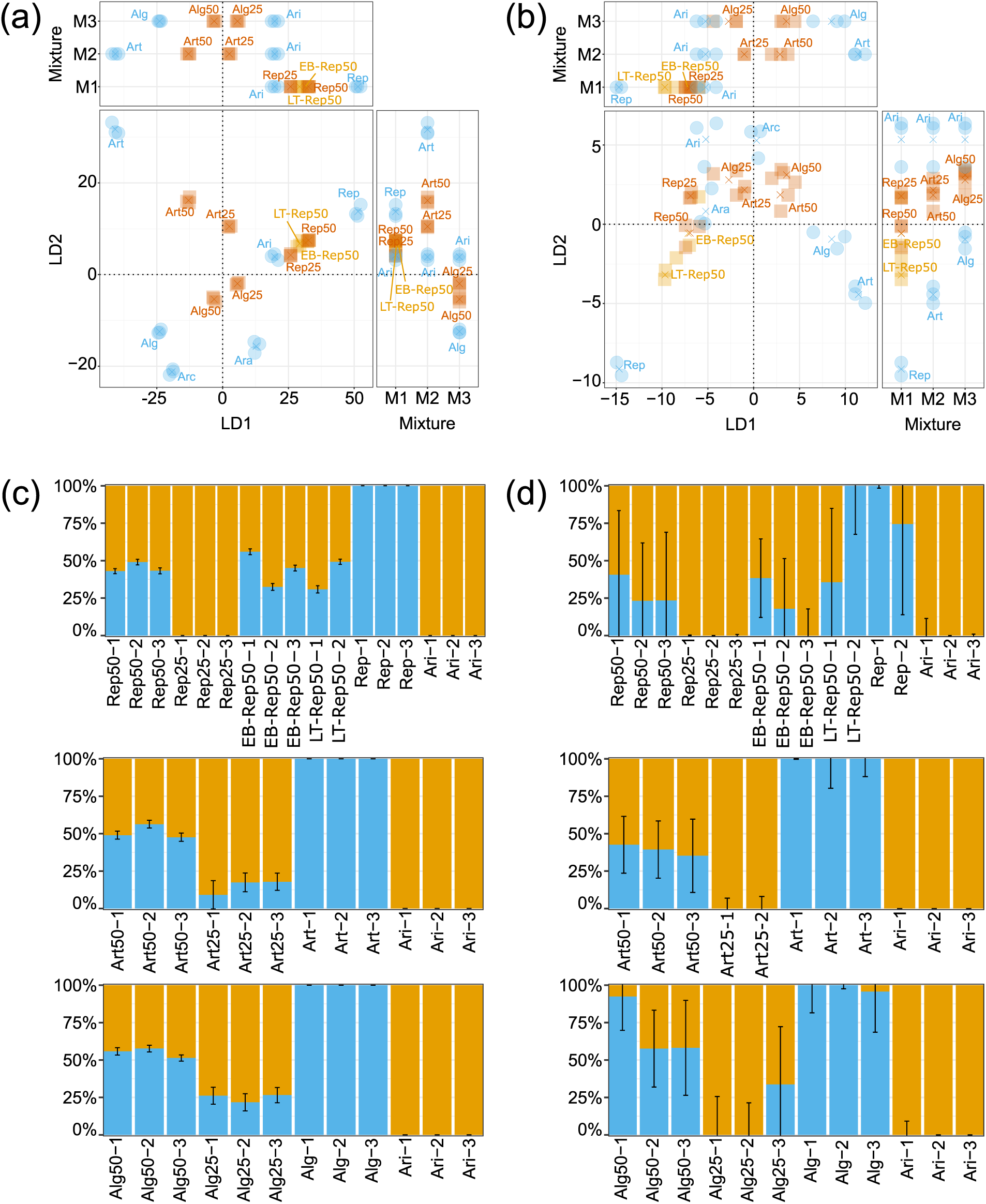
Discriminant analysis of principal components (a,b) and population structure (c,d) of an extended *Lolium perenne* L. sample set based on six cultivars: ‘Arara‘ (Ara), ‘Araias‘ (Ari), ‘Repentinia‘ (Rep), ‘Artonis‘ (Art), ‘Arcturus‘ (Arc), ‘Algira‘ (Alg). The sample set consists of single-cultivar samples (blue circles), mixed samples from a greenhouse (cultivar abbreviation followed by portion of cultivar in mixture; auburn squares), and mixed samples from a field experiments (sample names with experiment location as prefix (LT for Langenthal; EB for Eschenbach); beige squares) analysed using genotyping-by-sequencing (a) and multispecies amplicon sequencing (b). While Ari is present in all mixtures, Rep, Art, and Alg are present in M1, M2, and M3, respectively (Tab. 1). Replicate means are represented by crosses. Population structure analysis was based on genotyping-by-sequencing (c) and multispecies amplicon sequencing (d), with 95% confidence intervals. The probability of membership of each sample is indicated in beige (cluster of Ari) and blue. Sample names have the replicate number as suffix.

The population structure assessed separately for each mixture with ’ADMIXTURE’ were close to the expected ratios (Fig. 3(c)). The cultivars ‘Araias‘ and ‘Repentinia‘ were clearly separated and assigned to two separate clusters. Mixed samples containing the two cultivars at a 50:50-ratio (i.e., Rep50) were assigned to the two clusters at a similar ratio (i.e., slightly less than 50%). This was the case for the samples from both the greenhouse and the field experiment. Mixed samples with a 75:25-ratio (i.e., Rep25) were fully, but incorrectly assigned to the same cluster ‘Araias‘ belonged. The cultivars ‘Araias‘ and ‘Artonis‘ were clearly separated and assigned to two separate clusters. Mixed samples containing the two cultivars at a 50:50-ratio (i.e., Art50) were assigned to the two clusters at a similar ratio (i.e., approximately 50%). Mixed samples with a 75:25-ratio (i.e., Art25) were correctly assigned to the two clusters of the cultivars: While these samples were assigned to ‘Artonis‘ with a probability of approximately 25%, they were accordingly assigned to ‘Araias‘ with a probability of approximately 75%. The cultivars ‘Araias‘ and ‘Algira‘ were clearly separated and assigned to two separate clusters. Mixed samples containing the two cultivars at a 50:50-ratio (i.e., Alg50) and a 75:25-ratio (i.e., Alg25) were correctly assigned to the two clusters of the cultivars with similar separation patterns like the mixtures containing ‘Araias‘ and ‘Artonis‘. Results obtained with ’STRUCTURE’ were consistent with the ’ADMIXTURE’ analysis (Fig. S8a, S8b, S8c).

For the MSAS data, cultivars could be clearly separated using a DAPC (Fig. 3(b)). Based on the cross-validation procedure, four PCA axes were retained in the DAPC. The mixed samples containing cultivars from the greenhouse experiment at a 50:50-ratio were distinguished from and located in the middle between the corresponding cultivars. However, this differentiation pattern was only visible when LD2 was included either separately or together with LD1. For LD1, the differentiation pattern was not as expected for the mixed samples containing ‘Repentinia‘ or ‘Algira‘ (i.e., Rep50 and Rep25). For the samples from the field experiment, samples from Langenthal and Eschenbach exclusively showed an expected differentiation pattern using LD1 and LD2 separately, respectively. The differentiation pattern of the samples containing cultivars at a 75:25-ratio was closest to the expected pattern when both LD1 and LD2 were used. When used separately, LD1 and LD2 showed expected the differentiation patterns for mixed samples containing ‘Artonis‘ (i.e., Art25) and ‘Repentinia‘ (i.e., Rep25), respectively.

The population structure assessed separately for each mixture with ’ADMIXTURE’ were generally not congruent with the expected ratios (Fig. 3(d)). The cultivars ‘Araias‘ and ‘Repentinia‘ were separated and assigned to two separate clusters, with one exception. Mixed samples containing the two cultivars at a 50:50-ratio (i.e., Rep50) were incorrectly assigned to the two clusters of the cultivars: While these samples were assigned to ‘Repentinia‘ with a probability of approximately 25% instead of 50%, they were accordingly assigned to ‘Araias‘ with a probability of approximately 75% instead of 50%. Additionally, the variability between replicates was high, especially for the samples from the field experiment: While one replicate of EB-Rep50 was fully assigned to ‘Araias‘, one replicate of LT-Rep50 was fully assigned to ‘Repentinia‘. Mixed samples with a 75:25-ratio (i.e., Rep25) were fully, but incorrectly assigned to ‘Araias‘. The cultivars ‘Araias‘ and ‘Artonis‘ clearly separated and assigned to two separate clusters. Mixed samples containing the two cultivars at a 50:50-ratio (i.e., Art50) were slightly deviating from this ratio: While these samples were assigned to ‘Artonis‘ with a probability of approximately 40%, they were accordingly assigned to ‘Araias‘ with a probability of approximately 60%. Mixed samples with a 75:25-ratio (i.e., Art25) were fully, but incorrectly assigned to ‘Araias‘. The cultivars ‘Araias‘ and ‘Algira‘ separated and assigned to two separate clusters. Mixed samples containing the two cultivars at a 50:50-ratio (i.e., Alg50) deviated from this ratio: While two replicates were assigned to ‘Algira‘ and ‘Araias‘ with a probability of approximately 60% and 40%, respectively, one replicate was fully assigned to ‘Algira‘. For the samples with a 75:25-ratio (i.e., Alg25), two were fully assigned to ‘Araias‘ and one was assigned to ‘Algira‘ and ‘Araias‘ with a probability of approximately 30% and 70%, respectively. ’STRUCTURE’ and ’ADMIXTURE’ yielded comparable results (Fig. S8d, S8e, S8f).

## 4 Discussion

The methods, MSAS and GBS, were successfully used in this study. The workflows used, ranging from sampling to sample preparation and bioinformatics analysis, provide insight into their applicability for PGD monitoring of grassland.

PGD monitoring in grasslands starts with collecting reliable and representative samples. This is the most crucial, but also most time-consuming step. Increasing sampling efficiency and, therefore, decreasing time and labour would ease task allocation and would increase reliability of PGD monitoring studies as more samples could be collected (spacially and temporally). In this study, taking samples based on two parameters, number of individuals pooled in a sample and sampled leaf size, allowed highly reliable results to be obtained. For setting the pool sizes, the goal was to capture rare genotypes with a frequency of approximately 0.1. Therefore, we pooled 20 and 30 individuals per sample for the multispecies and the extended *L. perenne* to include at least one rare genotype with a frequency of 0.07 and 0.11 at a probability of 0.9, respectively (Crossa, 1989; Crossa and Vencovsky, 2011). Based on these pool sizes, we obtained highly reliable results confirmed by high correlations of allele frequencies between biological and technical replicates of *r>*0.9. For the leaf size, samples were collected based solely on leaf length. Therefore, other leaf characteristics, such as leaf width, were not taken into account, although leaf width differed even within species: tetraploid *L. perenne* cultivars had up to four times broader leaves than diploid cultivars. The results, however, were still reliable confirmed by the optimal separation pattern in mixtures of the extended *L. perenne* data set containing both diploid and tetraploid cultivars. In summary, this study contributes to more efficient PGD monitoring of grassland as samples containing only 20 to 30 individuals represented by solely leaf length provided reliable results.

Sampling for PGD monitoring is additionally time-consuming when it comes to multispecies grassland where, commonly, each species is sampled separately (e.g., Verwimp et al. 2018). In addition to the time constraint, botanical expertise is needed to separate species. Therefore, shifting species separation to another step in the PGD monitoring workflow could open up time-saving sampling methods, such as sampling from a pile of mown grassland species or from cow dung. In this study, we successfully shifted species separation from sampling to the computational part of the workflow. For this, we amplified MSAS target loci from different species present in a mixture and taxonomically assigned the sequences based on a reference database developed previously (Loera-Sánchez et al., 2022) and in this study. In general, the taxonomic assignment rate for legumes was higher than that for grasses, which is consistent with what has been reported for plastome DNA barcodes (Birch et al., 2017; Loera-Sánchez et al., 2020; Herzog and Latvis, 2021). For legumes, the high taxonomic assignment rates of above 70% explain the similar genetic differentiation patterns between accessions when comparing single-accession and multispecies samples. Even though the two legumes share similar life history (Ellison et al., 2006; Watson et al., 2000), the correct taxonomic assignment rate of *T. pratense* was higher and differed from that of *T. repens*. This could be explained by the allopolyploid nature of *T. repens* (Williams et al., 2012), leading to potentially higher divergence between the accessions used in this study and the reference MSAS sequence. For grasses, the lower taxonomic assignment rates explain the lower percentage of SNPs shared by the single- and multispecies seedling samples. Lower taxonomic assignment rates for grasses could be explained by their high genetic relatedness (Zhang et al., 2022) confirmed by evaluating the top incorrectly assigned species. For *F. pratensis* and *L. perenne*, these were closely related *L. perenne* and *L. multiflorum*, respectively (Cheng et al., 2016). For *D. glomerata*, however, this was *P. pratensis* which is more distantly related (Zhang et al., 2022). Additionally, the taxonomic assignment rate for *D. glomerata* was particularly lower than that of the other grasses. This could be explained by the high divergence between the candidate cultivar ’DG1525’ (i.e., cultivar A of *D. glomerata*) to the reference MSAS sequences of that species. In turn, this resulted in very low read mapping rates from multispecies samples to the *D. glomerata* reference sequences, so that no variant for *D. glomerata* was retained after filtering for minimum depth. Therefore, these results confirm potential difficulties in separating grassland species as the diversity within species tends to be higher than between species. Nevertheless, depending on the scope of PGD monitoring, our approach of shifting species separation to the more time-efficient computational part of PGD monitoring contributes to more efficient PGD monitoring. Therefore, our MSAS approach could be combined with even more time-efficient sampling techniques used to analyse environmental DNA, such as sampling airborne pollen (Van Haeften et al., 2024).

After sampling for PGD monitoring, the samples are processed, sequenced, and analysed. Depending on the scope of PGD monitoring, different methods can be chosen. An appropriate method should be (i) affordable, (ii) accurate, and (iii) scalable (Hoban et al., 2021a) where each method is a trade-off between these three properties. MSAS, which is a compromise of these three properties, was investigated in this study mainly with regard to its accuracy and scalability. Previously reported MSAS-based PGD metrics were limited to nucleotide diversity estimates in a single population. Here, we assessed the genetic differentiation among pairs of cultivars and ecotypes and observed that overall *F* _ST_ ranged from 0.04 to 0.06 (excluding *T. repens* and *D. glomerata*). Other studies assessing genetic differentiation between cultivars and ecotypes are scarce and are mainly available for *T. pratense* and *L. perenne*. For *T. pratense*, a study focused on Nordic cultivars and wild accessions and reported an overall *F* _ST_ = 0.032 based on ∼620 SNPs (Osterman et al., 2021). A *T. pratense* accession set of wider geographical origin (cultivars and European and Asian natural populations) showed an *F* _ST_ = 0.076 for ∼8000 SNPs (Jones et al., 2020). For *L. perenne*, a study grouped ecotypes and cultivars into clusters based on ∼500k SNPs (Blanco-Pastor et al., 2019). The authors reported an *F* _ST_ ranging from 0.015 to 0.065 when comparing clusters containing cultivars with clusters containing ecotypes. Therefore, the genetic differentiation estimates in our study fall within the same order of magnitude as those in other studies, which supports the accuracy of our approach. Furthermore, differentiation among accessions was significantly larger than within accessions and diversity estimates were highly precise across replicates. These results indicate that the nature of SNPs targeted by MSAS (i.e., SNPs located in coding regions and ultra-conserved-like elements, ULE) and the resulting SNP counts obtained, which are lower compared to other studies, are useful for estimating population differentiation.

Method selection for PGD monitoring is additionally dependent on the level of accuracy desired, not only among, but also within species. Therefore, methods suitable for genetically differentiating species might not be suitable for quantitatively assessing cultivar composition within a species (i.e., proportion of each cultivar within a mixture of cultivars). The latter type of analysis could be interesting along various steps during cultivar development. It could be applied in *in situ* conservation of grassland which provides the base material for breeding. Beside the quality of such protected areas, detection of potential shifts in the composition within species – either due to natural fluctuations or artificially (e.g., undesired overseeding activity) – is of high interest, but poorly studied (Wambugu and Henry, 2022). The method could also be applied to evaluate grassland restoration projects, where studies are scarce (Harźe et al., 2018). Beside breeding, the analysis type could be applied in variety testing. Potential variety shifts could be detected depending on various factors, such as management, which could serve as a basis for cultivars especially suited for specific managements, such as grazing. Additionally, that analysis type could be applied to assess cultivar purity (i.e., seeds only containing one cultivar) which is key for seed producers. However, these quantitative assessments of cultivar composition require methods of appropriate accuracy. In this study, we approximated the accuracy of MSAS in a multispecies and a single-species experiment. For the multispecies experiment, the suitability of MSAS in differentiating between accessions was evaluated. Exceptionally high genetic differentiation was detected between the candidate cultivar and the ecotype of *D. glomerata*. This high divergence might rely on the selection pressure in the candidate cultivar population during breeding, resulting in higher allele fixation compared to the ecotype (Espeland et al., 2017). Meanwhile, no genetic differentiation between the cultivar and the candidate cultivar of *T. repens* could be detected. While the divergence was expected to be lower compared to the other species where one of the two accessions was either an ecotype or a landrace, it was exceptionally lower than expected. This indicates that the genetic differentiation between both accessions falls below the detection limit of MSAS. The accuracy of MSAS was further assessed by preparing two samples where one contains accession A and the other accession B of each species. After further mixing these two mixed-species seedling samples at a 50:50-ratio, this resulting sample could be genetically differentiated from the two samples containing only accession A or B. Additionally, the mixing ratio could be displayed in a discriminant analysis of principal components (DAPC) where the 50:50-ratio sample was correctly located between the other two samples. The detection limit of MSAS was further evaluated by applying the method to a single-species experiment comprising an extended set of *L. perenne* samples of cultivars mixed at different ratios. Similarly to the mixed-species approach, the sample containing 50% of two *L. perenne* cultivars could be genetically differentiated from and were located between the two corresponding cultivars in a DAPC. However, samples containing 25% of one and 75% of the other cultivar did not segregate according to this ratio. Additionally, replicates showed high variability, which further indicates that MSAS reaches a detection limit for bi-cultivar samples containing less than 50% of a cultivar. Therefore, these results indicate limitations of MSAS being applicable to quantitatively assess cultivar composition.

Another method for PGD monitoring is GBS, a reduced genome representation method. This method produces genetic diversity estimates of high accuracy and could be applied to quantitatively assess cultivar compositions in PGD monitoring. To evaluate this application, the detection limit of GBS was evaluated and compared with that of MSAS using the same extended set of *L. perenne* samples. Similarly to MSAS, samples containing 50% of two *L. perenne* cultivars were located between the two corresponding cultivars in a DAPC. In addition to MSAS, samples containing 25% of one cultivar and 75% of ‘Araias‘ segregated according to this ratio and were located between the corresponding 50:50-sample and ‘Araias‘. Additionally, variability between replicates was low confirmed by high overlaps in the DAPC. The segregation pattern and the replicate variability could be confirmed by using two other methods, ADMIXTURE and STRUCTURE, where cluster memberships represented the actual cultivar ratios. These results indicate that GBS has a lower detection limit than MSAS and could detect cultivar shifts smaller than 25% in bi-cultivar samples. This quantitative separation of cultivar mixtures is a novel application of GBS as most other studies focused on differentiating cultivars without mixing them (e.g., Byrne et al. 2013). Verwimp et al. (2018), however, elaborated on the separation of cultivars in mixtures. The authors analysed, *inter alia*, changes in cultivar composition in mixtures containing two *L. perenne* cultivars initially sown at a 50:50-ratio. The allele frequencies of these mixtures were analysed across four years using a principal component analysis. As the samples came from a field experiment, the only reference on the ratio of the cultivars was the seed composition. Therefore, the authors could qualitatively track the temporal changes of cultivar mixtures, but could not further evaluate the quantitative detection limit of GBS. In the current study, the high accuracy of GBS was also confirmed by high genetic differentiation estimates between cultivars. Overall *F* _ST_ was 0.11, which was higher than that of MSAS (*F* _ST_ = 0.07) and similar to those of other studies (Blackmore et al., 2016; Blanco-Pastor et al., 2019). These high genetic differentiation estimates and accuracy of GBS enabled reconstructing the breeding history of the cultivars. Although all cultivars contain genetic material from the same Swiss ecotype collection, their breeding history differs. ‘Algira‘ and ‘Arcturus‘ are closely related as they originate from the same polycross (Grieder et al., 2015) resulting in the smallest *F* _ST_ values, the closest distance on the DAPC, and the high similarity in the ADMIXTURE analysis (K*<*6). ‘Arara‘ and ‘Araias‘ contain genetic material of a cultivar that is no longer on the list of recommended varieties, which resulted in smaller *F* _ST_ values and high similarity in the ADMIXTURE analysis (K*<*5). ‘Repentinia‘ and ‘Artonis‘ share the fewest genetic material with the other cultivars as they have the most polycrosses in their breeding history resulting in the highest *F* _ST_ values, large distances on the DAPC, and clear separation in the ADMIXTURE analysis (K*>*3). This is especially true for ‘Artonis‘ which is based on more distinct ecotypes collected at higher altitudes (Kempf et al., 2020) resulting in the highest separation in the ADMIXTURE analysis (K*>*2). Therefore, DAPC and ADMIXTURE could be successfully used to reconstruct the breeding history. However, the results of the methods slightly differed when analysing the cultivar composition. While the segregation pattern was similar for most samples, that of the mixed samples containing 75% of ‘Araias‘ and 25% of ‘Repentinia‘ differed between the methods as the ADMIXTURE algorithm did not separate these samples from the pure ‘Araias‘ samples. This is in agreement with similar studies (Tehseen et al., 2022; Deperi et al., 2018) and indicates the importance of including multiple methods to analyse genetic differentiation and approves this common practice (Miller et al., 2020).

We see great potential in MSAS and GBS being applied in PGD assessment of grassland. Both methods have strengths and limitations, making them applicable for different types of PGD analysis. MSAS could shift the labor-intensive species separation from the sampling to the computational part of the workflow. Additionally, the genetic diversity estimates were congruent across replicates and enabled analysing population differentiation of five grassland species at high accuracy. Nevertheless, MSAS reached a detection limit when quantitatively assessing cultivar compositions of *L. perenne*. The lower detection limit of GBS, however, allowed the separation of samples containing different cultivar compositions. Although this application was exclusively tested on *L. perenne* in the current study, GBS was also used to separate grassland species in another study (Wagemaker et al., 2021). Our results indicate that MSAS and GBS could be applied to a wide range of PGD monitoring studies. While both methods are cost-effective approaches (Hale et al., 2020; Carroll et al., 2018), MSAS is even more affordable than GBS. However, this comes at the expense of an underestimation of genetic diversity and a higher detection limit. Therefore, we anticipate that MSAS will be applied in PGD monitoring studies where resolution can be traded-off for a higher sample throughput. Meanwhile, we see the application of GBS in monitoring PGD at a smaller scale, but at higher accuracy. Beside these two approaches and with a further decrease in both the sequencing cost and the cost difference between sequencing platforms (Mustafa, 2024; Espinosa et al., 2024), approaches with even higher accuracy, such as whole-genome sequencing, might find their way to PGD monitoring. However, our study highlighted the complementing nature of MSAS and GBS equipping researchers with a flexible toolkit for PGD monitoring, suitable for various study scopes and resource levels.

## 5 Acknowledgements

We thank Christoph Grieder (Agroscope) for giving us access to the greenhouse to conduct the greenhouse experiments and providing information about breeding backgrounds of the plant material. We thank Hansjürg Fuhrimann, Hugo Jung, and Fabio Tanner for providing space to conduct and Daniel Suter (Agroscope) and Hansueli Hirschi (Agroscope) for establishing and maintaining the field experiments. We thank the Grassland Management and Ruminant Production Systems group of Beat Reidy (BFH-HAFL) for taking samples from the field experiments. We also thank Franco Widmer (Agroscope) and Tabea Koch (Agroscope) for giving us access to laboratory space and laboratory support, respectively. We thank Ingrid Stoffel-Studer (ETH Zurich) for supporting us in the laboratory and the Genetic Diversity Center (GDC; ETH Zurich) where we performed MSAS library preparation and sequencing. We thank the federal office for agriculture FOAG, InnoSuisse PSI, Swiss Grassland Society (AGFF), Eric Schweizer Ltd, Fenaco CP UFA-Samen, Otto Hauenstein Seeds Inc., Samen Steffen Ltd, and Delley seeds and plants Ltd for funding this project.

## Supporting Information A Materials and Methods

### A.1 Population structure analysis on the extended *L. perenne* samples

For the analysis using ’STRUCTURE’, VCFs were converted to ’STRUCTURE’ format using ’PGDSpider’ v2.1.1.5. (Lischer and Excoffier, 2012). In the resulting sample data files, a PopData column was added where the replicates of the same pure samples were assigned to the same population. The mixed samples were not assigned to a population (i.e., PopData value set to 0). For the groups containing both mixed and pure samples, two populations were assumed (i.e., MAXPOPS=2). The number of populations for the group containing only pure samples ranged from two to six (i.e., separate analysis for each number of populations). The ploidy, the burnin length, and the number of MCMC reps was set to diploid, 10,000, and 100,000, respectively. Additionally, admixture was assumed (i.e., NOADMIX=0), the marker type was set to co-dominant (i.e., RECESSIVEALLELES=0), and the PopData column was used (i.e., USEPOPINFO=1, except for the analysis of the pure sample group with MAXPOPS*<*6).

For the analysis using ’ADMIXTURE’, the same VCFs were used as for ’STRUCTURE’ and were converted to BED format using ’PLINK’ v1.9 (Purcell et al., 2007). The optimal number of clusters (K) was assessed using a 10-fold cross-validation procedure, 2000 bootstrap replicates were used to assess the standard error. The number of clusters ranged from two to the number of cultivar compositions in the group. Additionally, the multithreaded mode was used for the analysis of the GBS data to reduce computation time.

## Supporting Information B Figures and tables

**Fig. S1:**
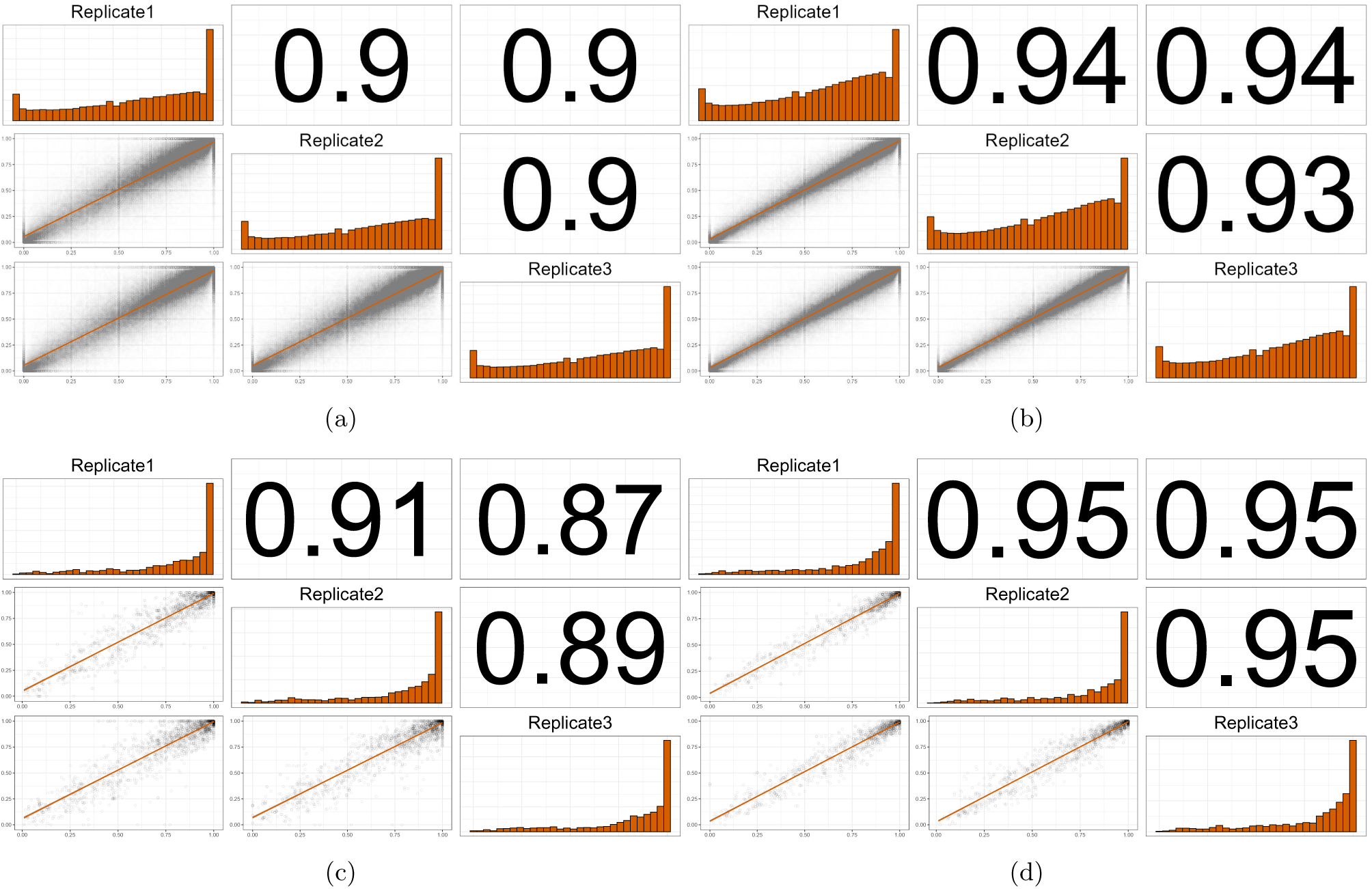
Comparison of allele frequencies between replicates of the extended *Lolium perenne* L. sample set of GBS (biological (a) and technical (b) replicates) and MSAS (biological (c) and technical (d) replicates). The upper triangle contains Spearman correlation coefficients, the diagonal contains the distribution of allele frequency values, and the lower diagonal contains the frequency of each allele summarised by linear regression lines.

**Fig. S2:**
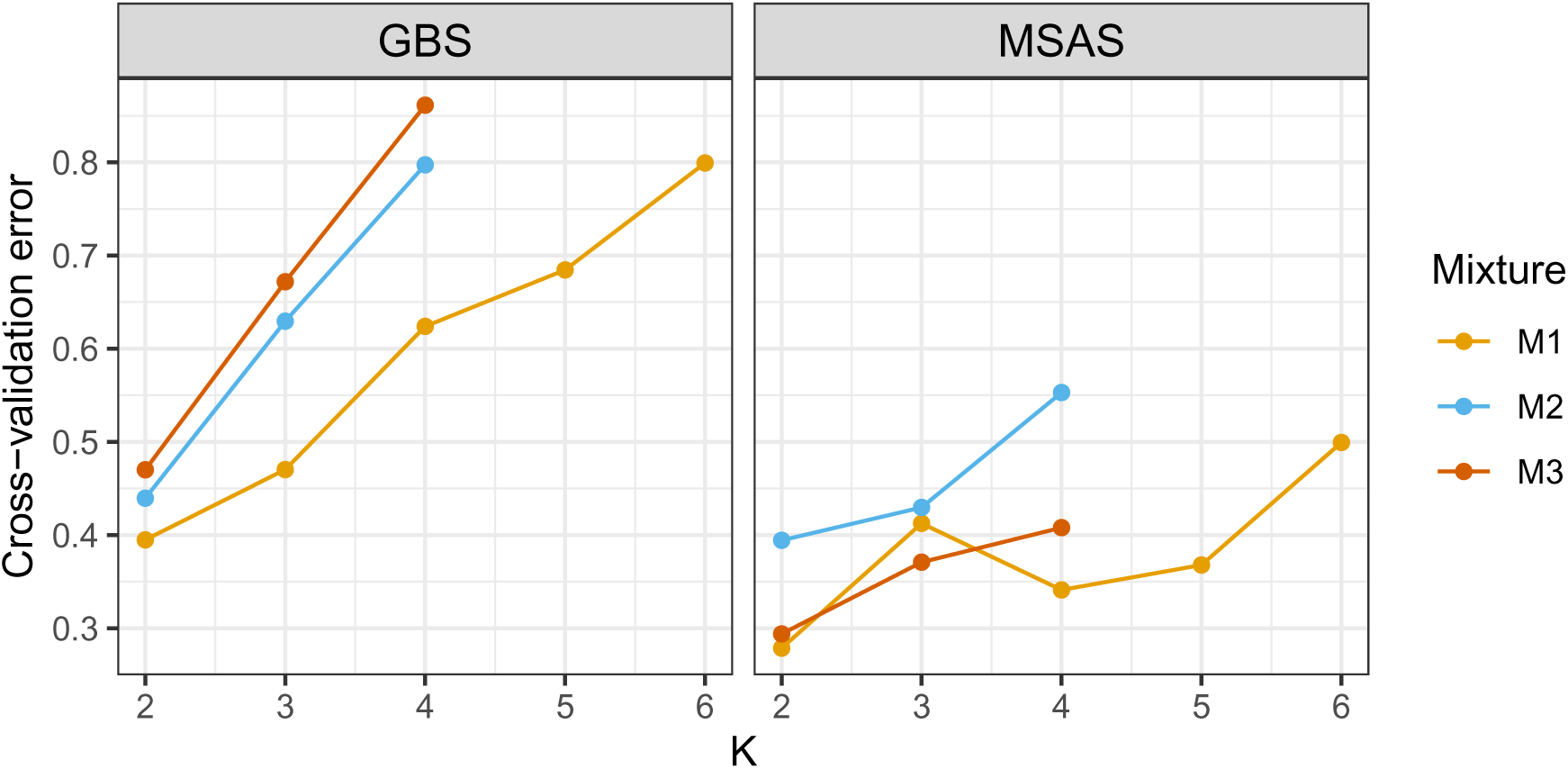
Cross-validation error per number of clusters (K) based on ’ADMIXTURE’ analysis of mixed samples of the extended *Lolium perenne* L. sample set generated with genotyping-by-sequencing (GBS) and multispecies amplicon sequencing (MSAS). ‘Araias‘, which is present in all mixed samples, is mixed with ‘Repentinia‘ (M1; contains 6 populations based on samples from the greenhouse and the field experiment), ‘Artonis‘ (M2; contains 4 populations based on samples from the greenhouse), and ‘Algira‘ (M3; contains 4 populations based on samples from the greenhouse).

**Fig. S3:**
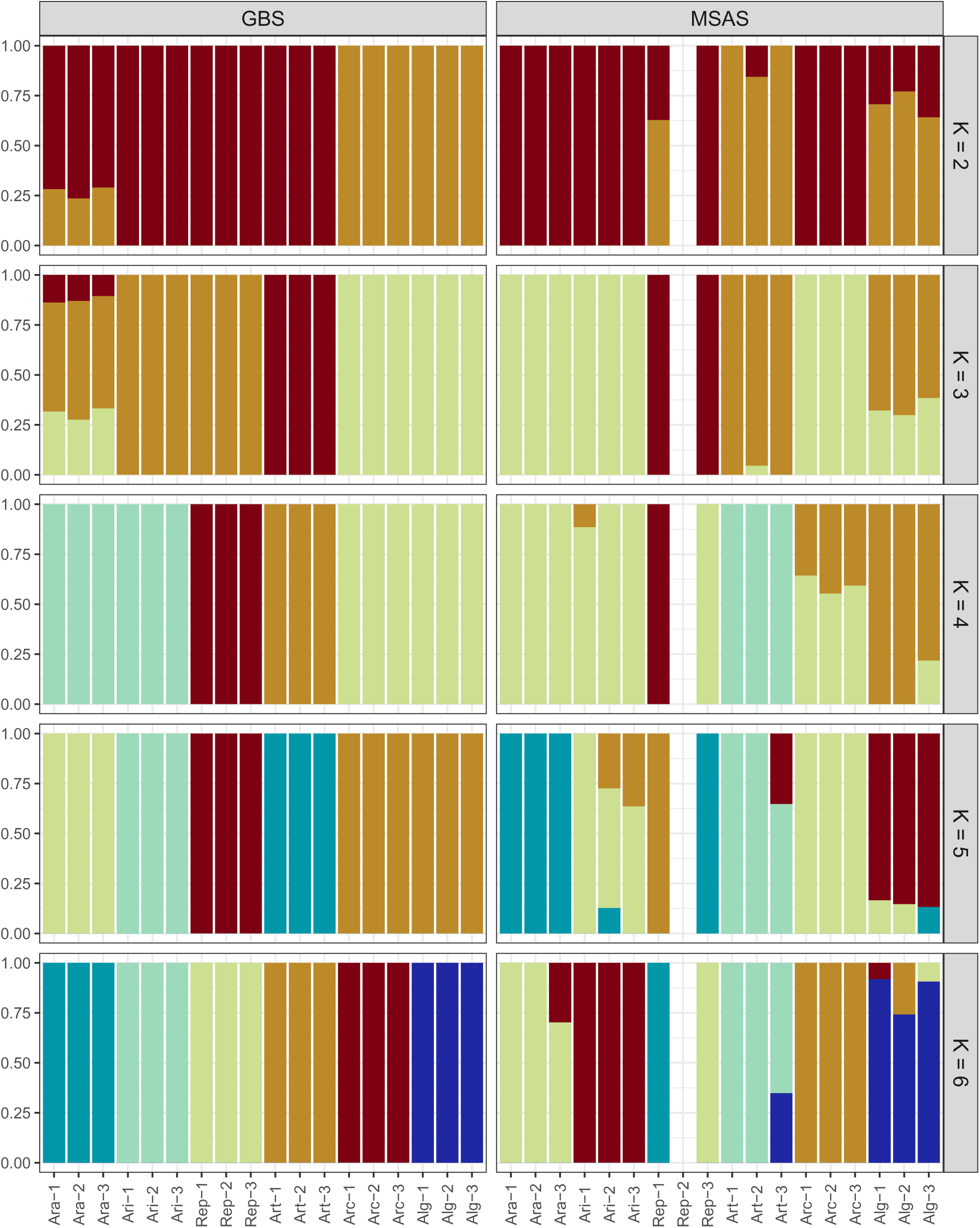
Population structure (2≤ K≤ 6) based on ’ADMIXTURE’ analysis of pure samples of extended *Lolium perenne* L. sample set generated with genotyping-by-sequencing (GBS) and multispecies amplicon sequencing (MSAS). The cultivar abbreviation is followed by replicate number.

**Fig. S4:**
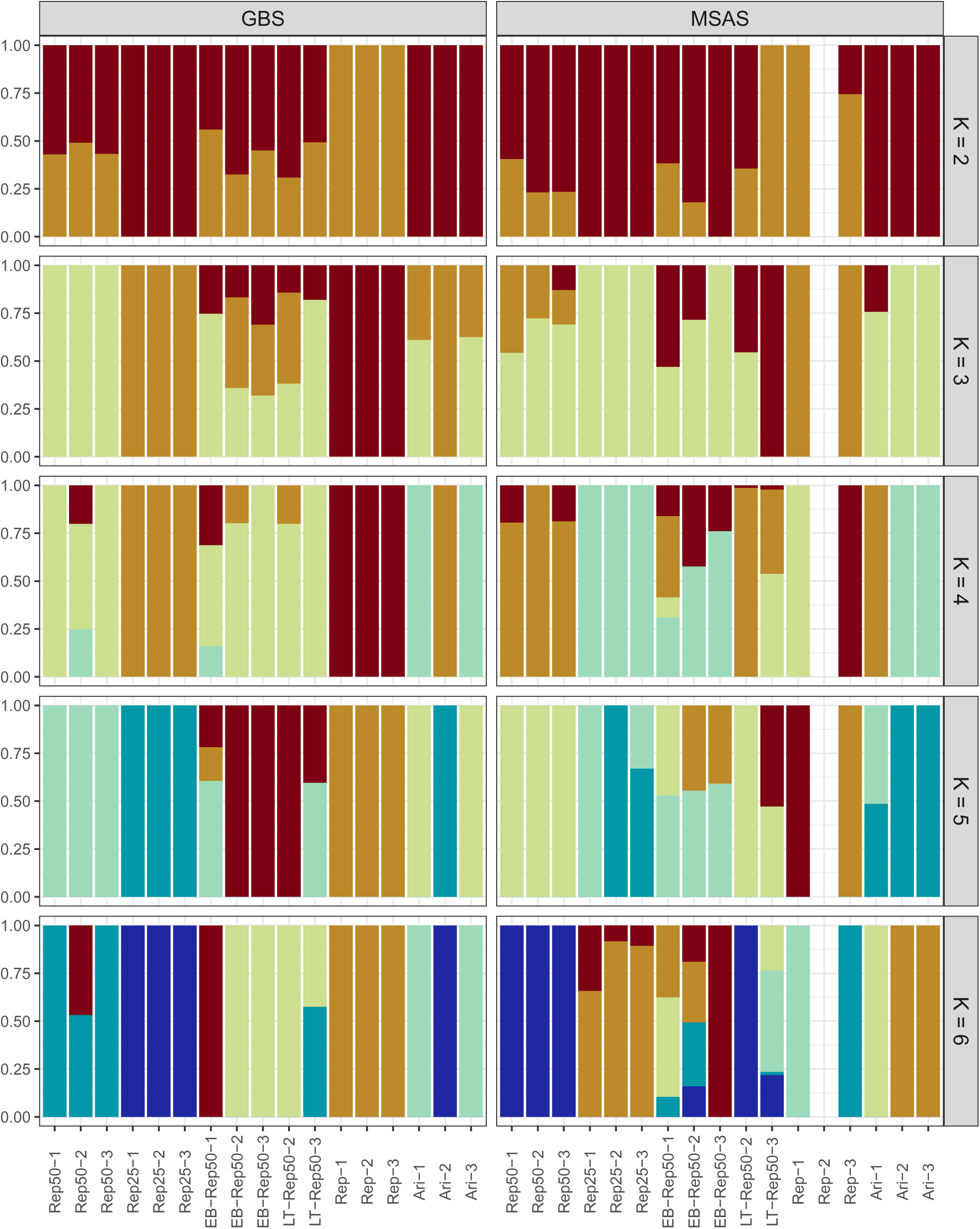
Population structure (2≤ K≤ 6) based on ’ADMIXTURE’ analysis of mixed samples of extended *Lolium perenne* L. sample set generated with genotyping-by-sequencing (GBS) and multispecies amplicon sequencing (MSAS). ‘Araias‘ (Ari) is mixed with ‘Repentinia‘ (Rep). The cultivar abbreviation is followed by the proportion of a cultivar in a mixture (e.g., Rep25 contains 25% Rep) and the replicate number. Samples from the field experiment correspond to Rep50 and were sown at two locations (EB and LT as sample name prefix).

**Fig. S5:**
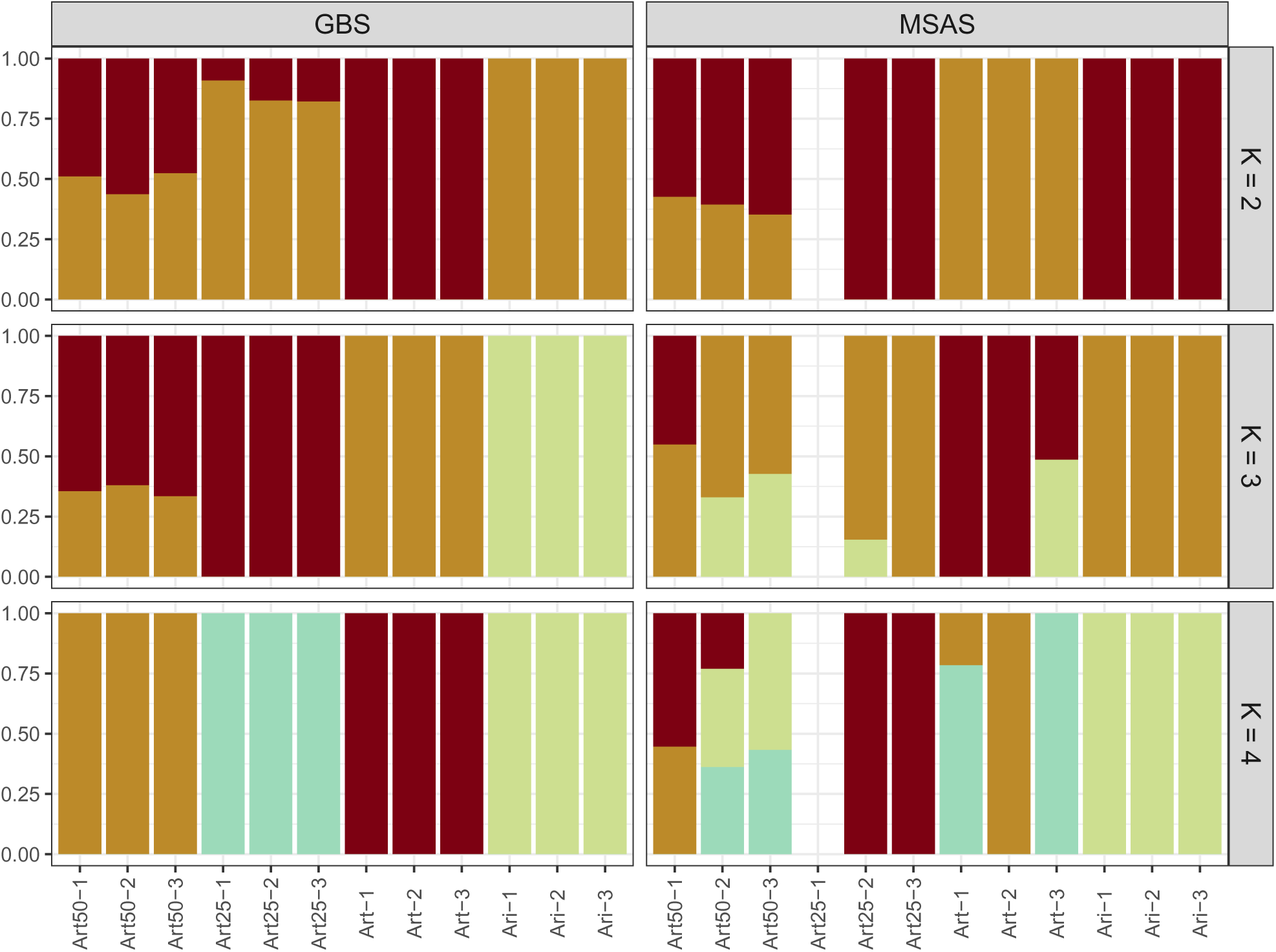
Population structure (2≤ K≤ 4) based on ’ADMIXTURE’ analysis of mixed samples of extended *Lolium perenne* L. sample set generated with genotyping-by-sequencing (GBS) and multispecies amplicon sequencing (MSAS). ‘Araias‘ (Ari) is mixed with ‘Artonis‘ (Art). The cultivar abbreviation is followed by the proportion of a cultivar in a mixture (e.g., Art25 contains 25% Art) and the replicate number.

**Fig. S6:**
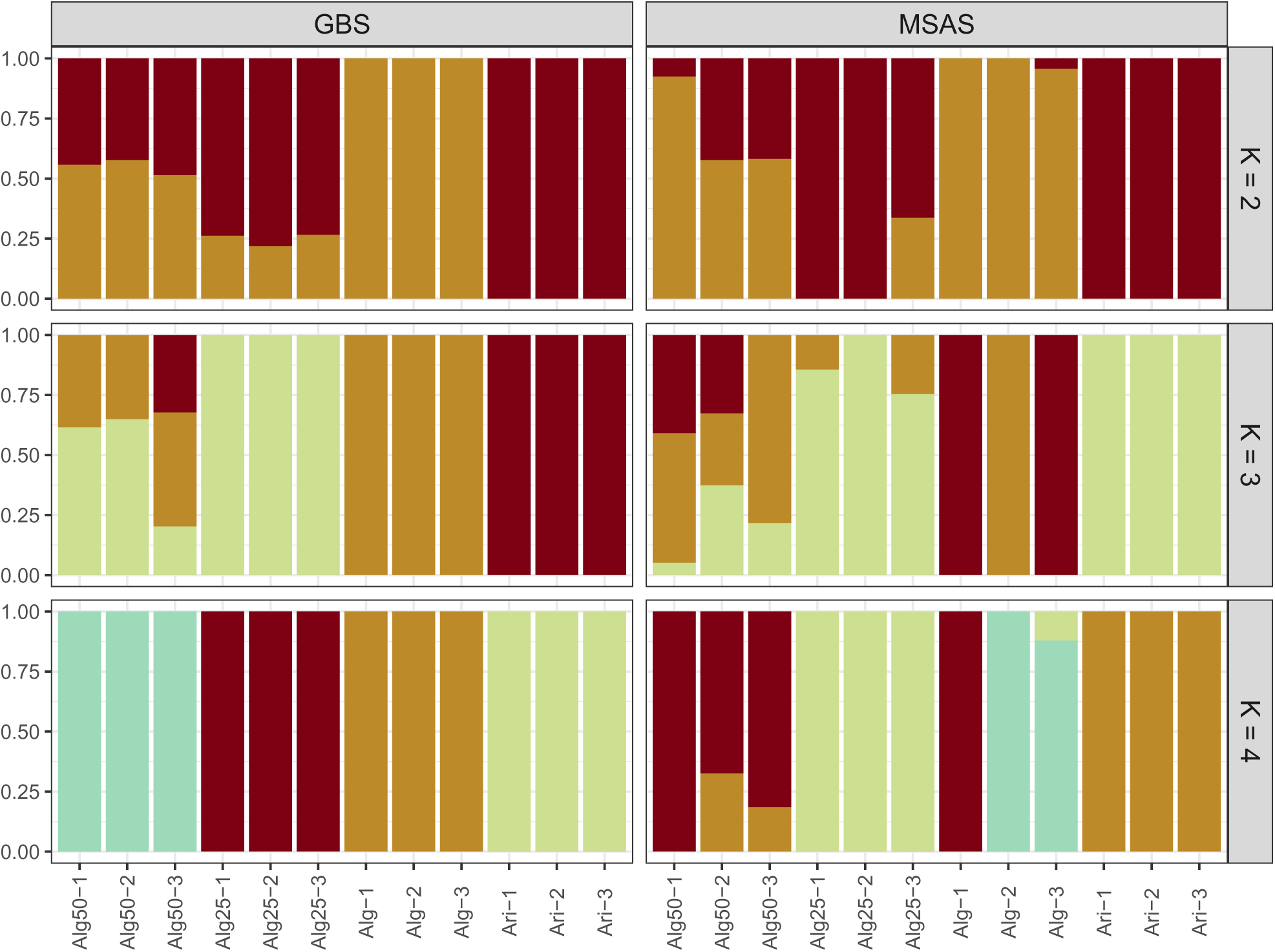
Population structure (2≤ K≤ 4) based on ’ADMIXTURE’ analysis of mixed samples of extended *Lolium perenne* L. sample set generated with genotyping-by-sequencing (GBS) and multispecies amplicon sequencing (MSAS). ‘Araias‘ (Ari) is mixed with ‘Algira‘ (Alg). The cultivar abbreviation is followed by the proportion of a cultivar in a mixture (e.g., Alg25 contains 25% Alg) and the replicate number.

**Fig. S7:**
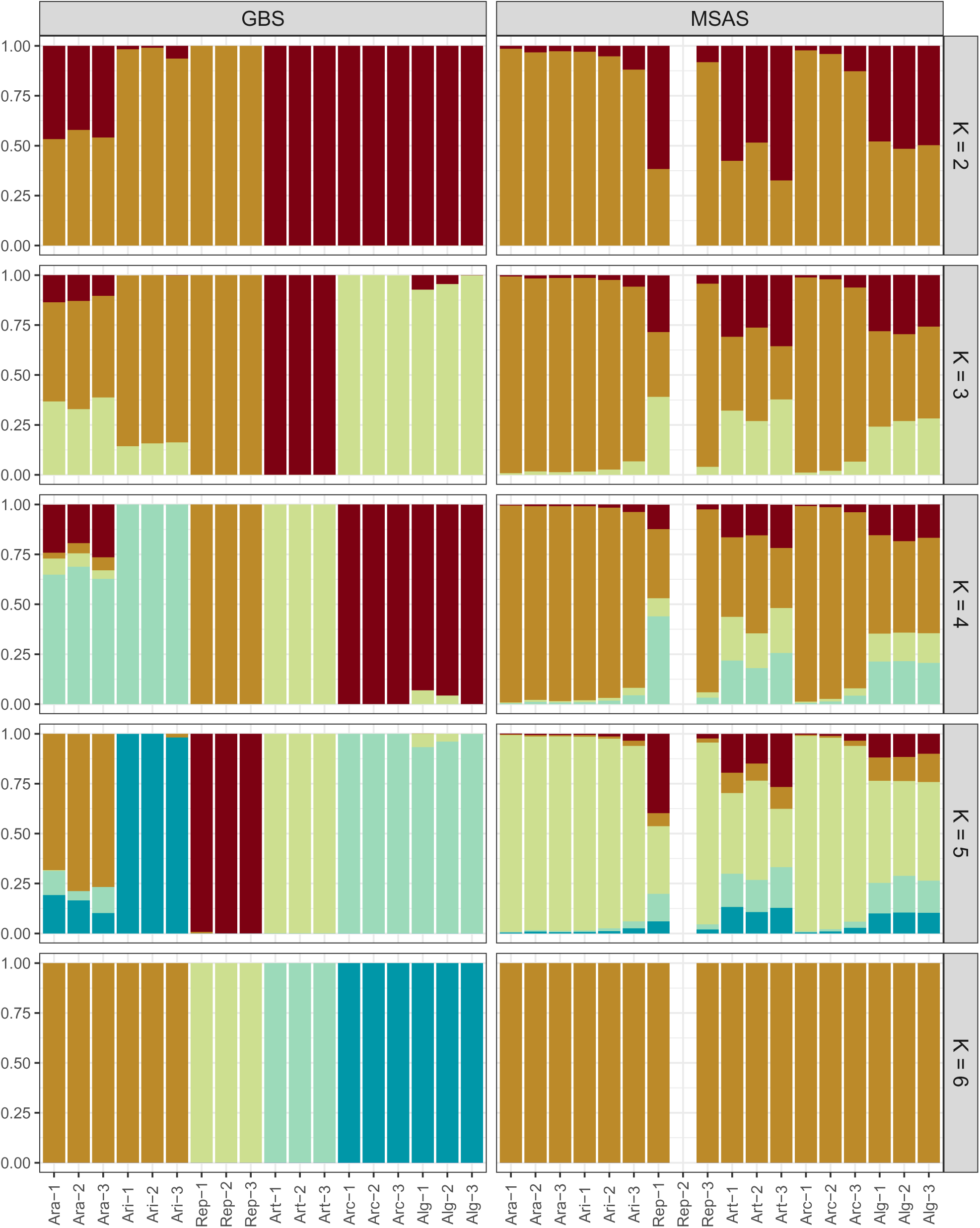
Population structure (2≤ K≤ 6) based on ’STRUCTURE’ analysis of pure samples of extended *Lolium perenne* L. sample set generated with genotyping-by-sequencing (GBS) and multispecies amplicon sequencing (MSAS). The cultivar abbreviation is followed by replicate number.

**Fig. S8:**
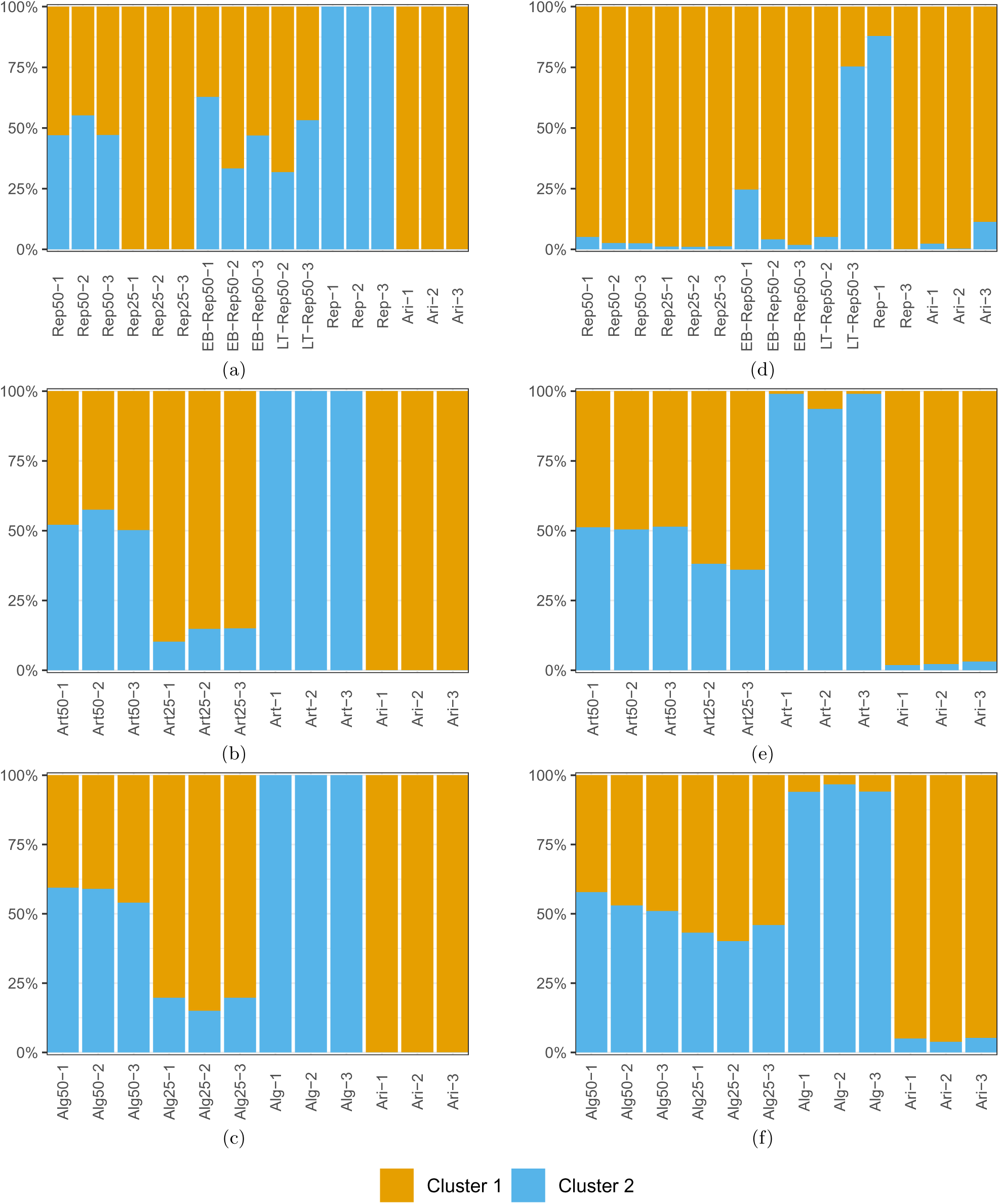
Population structure based on ’STRUCTURE’ analysis of the mixed samples of the extended *Lolium perenne* L. sample set generated with genotyping-by-sequencing (a-c) and multispecies amplicon sequencing (d-f). The probability of membership of each sample is indicated in beige (Cluster 1) and blue (Cluster 2). ‘Araias‘ (Ari), which is present in all mixed samples, is mixed with ‘Repentinia‘ (Rep; a,d), ‘Artonis‘ (Art; b,e), and ‘Algira‘ (Alg; c,f). For the mixed samples, the cultivar abbreviation is followed by the proportion of a cultivar in a mixture (e.g., Rep25 contains 25% Rep) and the replicate number. Samples from the field experiment correspond to Rep50 and were sown at two locations (EB and LT as sample name prefix).

**Table S1:**
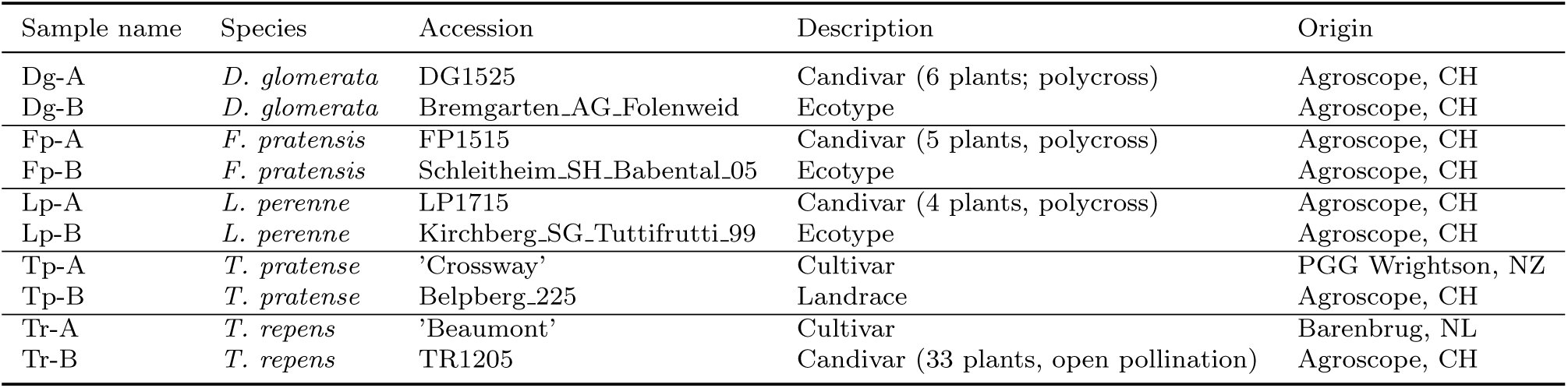
Accession compositions of the single-accession seedling samples.

**Table S2:** Accession compositions of the mixed-species seedling samples.

| Sample name | Composition of DNA mixture |
| --- | --- |
| MS-A100 | 20% Dg-A |
|  | 20% Lp-A |
|  | 20% Fp-A |
|  | 20% Tp-A |
|  | 20% Tr-A |
| MS-B100 | 20% Dg-B |
|  | 20% Lp-B |
|  | 20% Fp-B |
|  | 20% Tp-B |
|  | 20% Tr-B |
| MS-AB50 | 50% MS-A100 |
|  | 50% MS-B100 |

**Table S3:**
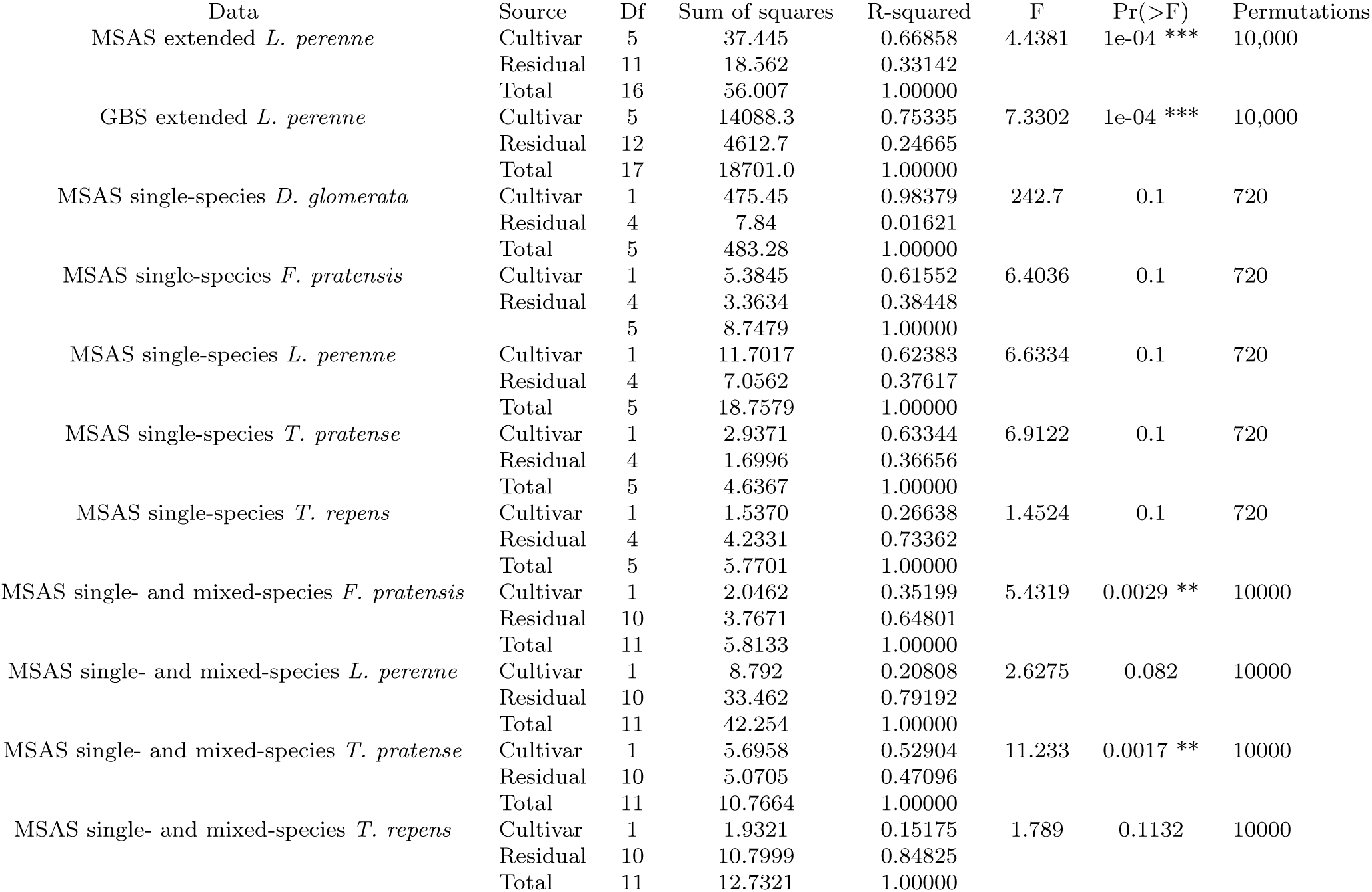
PERMANOVA results.

**Table S4:** Orthologues of MSAS loci in two reference genomes.

| Locus | *Brachypodium distachyon* v3.0 | *Arabidopsis thaliana* TAIR10 |
| --- | --- | --- |
| ortho-283 | BRADI_1g32610v3 | AT3G45100 |
| ortho-1542 | BRADI_3g30847v3 | AT5G53580 |
| ortho-1455 | BRADI_1g20730v3 | AT1G77930 |
| ortho-473 | BRADI_2g41580v3 | AT1G17470,AT1G72660 |
| uce-11005138 | BRADI_1g63540v3 | AT5G38460 |
| ortho-1288 | BRADI_1g63540v3 | AT5G38460 |
| uce-11005009 | BRADI_1g73110v3 | AT1G80680 |
| uce-11005008 | BRADI_1g73110v3 | AT1G80680 |
| ortho-1375 | BRADI_4g02070v3 | AT1G16650 |
| ortho-165 | BRADI_4g31680v3 | AT1G50480 |
| uce-11004158 | BRADI_3g02900v3 | AT1G70570 |
| ortho-1186 | BRADI_3g43450v3 | AT3G25410 |
| uce-11004715 | BRADI_3g60040v3 | AT2G22530 |
| ortho-1347 | BRADI_3g60150v3 | AT2G44020 |
| uce-11004502 | BRADI_1g35670v3 | AT5G42390 |
| ortho-1019 | BRADI_1g44820v3 | AT1G10520 |
| ortho-1245 | BRADI_3g14530v3 | AT4G15240 |
| ortho-1110 | BRADI_1g41970v3 | AT3G20440 |
| ortho-1318 | BRADI_1g31400v3 | AT5G04050 |

